# Multiomic and Spatial Profiling of Colorectal Tissue Reveals Viral Persistence and Immune Dysregulation in Long COVID

**DOI:** 10.64898/2026.08.07.743616

**Authors:** Brian LaFranchi, David P. Maison, Joanna Vinden, Antonio E. Rodriguez, Ali Tout, Lilian Grimbert, Emanuel Velazquez, Urmila Vudali, Belen Altamirano Poblano, Thomas Dalhuisen, Javier Cattle, Tony R. Figueroa, Yuki Fudotan, Michael Luna, Dylan Ryder, Monika Deswal, Brent S. Abel, James Lynch, Amanda Lipford, Nabeel Razi, Carly B. Steifman, Henry N. McCann, Nick Kataria, Valerie Girling, Reuben Thomas, Chao Wang, Amelia N. Deitchman, Shreya Patel, Michela Traglia, Zian H. Tseng, Gyula Szabo, Zoltan Laszik, Alex Farrow, Nicolaas Zwart, Nanami Sumimoto, Venice Servellita, Rebecca Hoh, Emily A. Fehrman, J. Daniel Kelly, Jeffrey N. Martin, Steven G. Deeks, Charles Y. Chiu, Ma Somsouk, Michael J. Peluso, Timothy J. Henrich

## Abstract

Long COVID (LC) – a chronic condition characterized by persistent, debilitating symptoms following SARS-CoV-2 infection – has emerged as a major public health challenge. Although many interrelated mechanisms have been proposed as drivers of LC, the root causes have yet to be identified, posing significant challenges for therapeutic development. While many blood-based studies have been conducted, they have not yielded conclusive mechanistic insights into LC pathogenesis. Attention has therefore turned toward direct tissue investigation, with the gastrointestinal (GI) tract becoming a major focus due to evidence that virus or viral components can persist at this site for months to years following an episode of COVID-19. Here, we performed a high-dimensional characterization of colorectal tissue and peripheral blood in a highly characterized cohort of 44 people with LC and 13 recovered controls. We profiled SARS-CoV-2 persistence, host immune responses, and tissue inflammation using bulk and single-cell RNA sequencing, nCounter RNA probe hybridization, quantitative PCR, metagenomic next-generation sequencing, plasma proteomics, high-dimensional spectral flow cytometry, in situ-hybridization/immunohistochemistry, and single-cell digital spatial omics. Our results support a model in which LC is driven by long-term immune dysregulation and perturbations of the regulatory gut immune environment which imply ongoing viral persistence, although direct viral detection was only observed in a subset of participants. Specifically, we identify a tissue-based transcriptional environment in which SARS-CoV-2 activates innate myeloid immune signaling, driving chronic inflammation while simultaneously downregulating pathways responsible for immune-mediated clearance of infected cells, including antigen presentation, phagocytosis, cytotoxic immune cell trafficking, and granzyme production. Importantly, signatures in peripheral blood are considerably weaker than those observed in tissue. Together, these findings provide a direct biological rationale for therapeutic strategies in LC aimed at enhancing or redirecting cytotoxic immune function to overcome immune dysregulation and clear persistent viral reservoirs.

## INTRODUCTION

Long COVID (LC), a chronic condition characterized by persistent, debilitating symptoms for months to years following SARS-CoV-2 infection^1^, can lead to significant impairment in daily functioning and quality of life^2^. Multiple mechanisms, including SARS-CoV-2 persistence, autoimmunity, vascular inflammation, gut dysbiosis, and immune dysregulation, have been proposed as drivers of the condition^3^. However, the biology of LC remains incompletely understood^3^. This has hampered the design of clinical trials and the identification of potential therapeutics for the millions of individuals worldwide affected by the condition^4,5^.

Most studies of LC have relied on measurements in easily accessible compartments such as peripheral blood or stool, which may not reflect the tissue-level processes driving the condition^6^. Direct tissue-based investigation has therefore been proposed as critical to understanding the underlying biology of LC^4^. The gastrointestinal (GI) tract has become a major focus of this effort, given evidence that virus or viral components can persist in this compartment for months to years following an episode of COVID-19^6–11^. For example, we previously reported the presence of single- and double-stranded SARS-CoV-2 RNA in rectosigmoid tissue for two years following initial infection, in the absence of known interval reinfection^12^.

Others have similarly reported viral RNA persistence in colorectal tissue months after infection^6–11^, including in people with inflammatory bowel disease^12^. However, tissue-based studies to date have been limited in size and scope, with small numbers of participants, confinement to specific phases of the pandemic, or a lack of comparable control groups of individuals who recovered following COVID-19^4^.

Here, we report findings from a highly characterized cohort of 44 people with LC and 13 recovered controls, longitudinally evaluated over a median of two years following initial SARS-CoV-2 infection across variant eras, in whom we performed deep, high-dimensional profiling of rectosigmoid tissue and peripheral blood using a combination of single-molecule, single-cell, and multiomic approaches. Overall, our findings reveal SARS-CoV-2 persistence in colorectal tissue predominantly but not exclusively in people with LC, accompanied by profound immune dysregulation in this tissue compartment that distinguishes LC from recovered individuals. Specifically, we identify a transcriptional environment in which SARS-CoV-2 – whether through direct viral infection or bystander viral effects – activates innate myeloid immune signaling, driving chronic inflammation while simultaneously downregulating pathways responsible for immune-mediated clearance of infected cells, including antigen presentation, phagocytosis, cytotoxic immune cell trafficking, and granzyme production. Importantly, blood signatures are considerably weaker than those observed in tissue, and SARS-CoV-2 detection in blood was rare. Collectively, these findings shed new light on the biological underpinnings of LC in tissue, with direct implications for the design of mechanism-based therapeutic trials for the condition.

## RESULTS

### Clinical cohort

Rectosigmoid biopsies and peripheral blood were collected from 57 participants enrolled in the UCSF Long-term Impact of Infection with Novel Coronavirus (LIINC) program between October 2020 and May 2025, including 44 individuals with LC and 13 recovered controls.

Demographic characteristics, clinical features, and underlying medical conditions are summarized in **Table 1**. The median age of the cohort was 45 years (IQR 38–57), 61% of participants were female, and the cohort was predominantly White (74%), with Asian (11%), Hispanic/Latino (11%), and Black/African American (3.5%) participants also represented. The most common comorbidities were pre-COVID-19 lung disease (i.e., asthma or COPD; 21%) and hypertension (14%). Notably, only four participants – all with LC – had required hospitalization during acute COVID-19. At the time of gut biopsy, a median of 689 days had elapsed since initial SARS-CoV-2 infection and a median of 536 days since the most recent known infection. In 30% of the cohort, the index infection occurred in the pre-Omicron era (ancestral through Delta variants, March 2020 to November 2021); in the remainder of cases, the index infection occurred during the Omicron era (after December 2021). The majority of participants (74%) had experienced their first episode of COVID-19 following completion of the SARS-CoV-2 primary vaccination series; all but two participants (both in the LC group) had completed the primary COVID-19 vaccination series prior to biopsy.

**Table 1:** Participant demographics and clinical characteristics.

| Characteristic | Overall<br>N = 57<sup>1</sup> | Long COVID<br>N = 44<sup>1</sup> | Recovered<br>N = 13<sup>1</sup> | p-value |
| --- | --- | --- | --- | --- |
| **Age** (years) | 45 (38, 57) | 45 (38, 56) | 39 (33, 58) | 0.8 |
| **Sex** |  |  |  | 0.5 |
| Female | 35 (61%) | 26 (59%) | 9 (69%) |  |
| Male | 22 (39%) | 18 (41%) | 4 (31%) |  |
| **Race/Ethnicity** |  |  |  | 0.8 |
| American Indian or Alaska Native | 1 (1.8%) | 1 (2.3%) | 0 (0%) |  |
| Asian | 6 (11%) | 5 (11%) | 1 (7.7%) |  |
| Black/African American | 2 (3.5%) | 1 (2.3%) | 1 (7.7%) |  |
| Hispanic/Latino | 6 (11%) | 5 (11%) | 1 (7.7%) |  |
| White | 42 (74%) | 32 (73%) | 10 (77%) |  |
| **Medical comorbidities** |  |  |  |  |
| Autoimmune disease | 7 (12%) | 5 (11%) | 2 (15%) | 0.7 |
| Cancer<sup>2</sup> | 1 (1.8%) | 0 (0%) | 1 (7.7%) | 0.2 |
| Diabetes | 2 (3.5%) | 2 (4.5%) | 0 (0%) | >0.9 |
| Hypertension | 8 (14%) | 8 (18%) | 0 (0%) | 0.2 |
| Lung disease<sup>3</sup> | 12 (21%) | 11 (25%) | 1 (7.7%) | 0.3 |
| **Body mass index** (kg/m<sup>2</sup>) |  |  |  | 0.9 |
| <25 | 29 (52%) | 23 (53%) | 6 (46%) |  |
| 25-29.9 | 18 (32%) | 13 (30%) | 5 (38%) |  |
| >= 30 | 10 (18%) | 8 (18%) | 2 (15%) |  |
| **Hospitalized for COVID-19** | 4 (7.0%) | 4 (9.1%) | 0 (0%) | 0.6 |
| **Antiviral treatment for acute COVID-19<sup>4</sup>** | 9 (16%) | 7 (16%) | 2 (15%) | >0.9 |
| **First SVC2 infection era** |  |  |  | 0.2 |
| March 2020 – April 2021 (ancestral) | 14 (25%) | 13 (30%) | 1 (8%) |  |
| May 2021 – November 2021 (Delta) | 3 (5.3%) | 3 (6.8%) | 0 (0%) |  |
| >December 2021 (Omicron) | 40 (70%) | 28 (64%) | 12 (92%) |  |
| **Time since first SCV2 infection** (days) | 689 (500, 863) | 725 (501, 878) | 636 (487, 729) | 0.4 |
| **Time since most recent SCV2 infection** (days) | 536 (361, 764) | 511 (315, 778) | 636 (487, 729) | 0.3 |
| **Vaccinated prior to first SCV2 infection** | 42 (74%) | 30 (68%) | 12 (92%) | 0.15 |
| **Vaccinated prior to colorectal biopsy<sup>5</sup>** | 55 (96%) | 42 (95%) | 13 (100%) | >0.9 |
| **Quality of life by visual analog scale<sup>5,6</sup>** | 65 (50, 75) | 60 (49, 70) | 82 (80, 85) | <0.001 |
| **Long COVID symptom count<sup>5</sup>** | - | 8 (5, 10) | - | - |
| **Symptoms reported<sup>7</sup>** |  |  |  |  |
| Fatigue | - | 39 (89%) | - |  |
| Cardiopulmonary | - | 18 (41%) | - |  |
| Gastrointestinal | - | 21 (48%) | - |  |
| Smell/Taste disturbance | - | 7 (16%) | - |  |
| Neurocognitive | - | 41 (93%) | - |  |
| Post-Exertional Malaise | - | 29 (66%) | - |  |
| Sleep disturbance | - | 27 (61%) | - |  |

<sup>1</sup>n (%); Median (IQR). <sup>2</sup>Cancer requiring treatment within the 2 years before COVID-19. <sup>3</sup>Asthma, COPD, emphysema or bronchitis experienced in the 5 years prior to COVID-19. <sup>4</sup>nirmatrelvir/ritonavir, molnupiravir, or remdesivir. <sup>5</sup>At time of biopsy. <sup>6</sup>Data missing for one participant. <sup>7</sup>**Cardiopulmonary:** cough, shortness of breath, heart palpitations, chest pain. **Gastrointestinal:** diarrhea, nausea, loss of appetite, abdominal pain, vomiting, constipation. **Neurocognitive:** concentration problems, dizziness, balance problems, neuropathy, vision problems.

As part of the parent cohort procedures, participants completed structured, interviewer-administered questionnaires to characterize the presence and severity of symptoms that newly emerged or worsened following SARS-CoV-2 infection. This enabled identification of participants meeting clinical criteria for LC consistent with the widely accepted case definitions put forth by the World Health Organization (WHO)^13^ and the National Academies of Sciences, Engineering, and Medicine (NASEM)^1^, as well as their classification of LC into clinical domains. Among participants who met the case definition of LC, the most prevalent symptom domains were neurocognitive impairment (93%), fatigue (89%), post-exertional malaise (PEM; 66%), and sleep disturbance (61%); cardiopulmonary and gastrointestinal symptoms were reported by 41% and 48%, respectively. Symptom domains were not mutually exclusive. Participants with LC reported a median of 8 symptoms (IQR 5-10) at the time of sampling. The control group had recovered following SARS-CoV-2 infection and reported no residual symptoms. Overall health on a 100-point visual analog scale (VAS) was significantly lower in the LC group compared to the recovered control group (median 60 vs 82, p<0.001).

Participants were followed longitudinally before and after colorectal tissue sampling by flexible sigmoidoscopy. Suspected or confirmed SARS-CoV-2 reinfections were systematically documented based on participant-reported symptoms and home or study-administered SARS-CoV-2 testing. In most cases (45/57, 78.9%), nasal SARS-CoV-2 PCR testing was performed on the day of the gut biopsy (n=32) or within 14 days before or after the procedure (n=13) to exclude occult or asymptomatic reinfection. In the 12 cases preceding implementation of this systematic testing procedure, participants were screened to exclude intervening symptomatic illness or known COVID-19 exposures during the preceding 30 days.

#### SARS-CoV-2 detection in colorectal tissue

To assess the presence, frequency, and tissue distribution of persistent SARS-CoV-2, we applied a complementary set of sequencing-based, proteomic, and spatial detection methods to colorectal tissue. To help define the specificity of tissue-based assays, we incorporated orthogonal viral detection and quantitation assays. In addition, our analyses included tissue specimens collected prior to the emergence of the SARS-CoV-2 pandemic as experimental controls to better define assay specificity.

First, using 5 µm thick formalin-fixed, paraffin embedded (FFPE) tissue sections, we performed multiplexed in-situ hybridization (ISH) and immunohistochemical (IHC) staining to detect SARS-CoV-2 single-stranded Spike and double-stranded ORF1A/B RNA, as well as SARS-CoV-2 Spike protein. We then co-extracted cellular RNA and DNA from cryopreserved bulk colorectal tissues and performed (1) unbiased metagenomic next-generation sequencing (mNGS) with and without enrichment using a viral capture probe panel (Twist Bioscience, South San Francisco, CA) targeting more than 1,000 eukaryotic viruses, including known human pathogens (2) multiplexed RNA probe hybridization quantitation (nCounter, nanoString Technologies, Seattle, WA) incorporating 807 human host-viral response transcripts and viral transcripts across multiple genomic regions of SARS-CoV-2, SARS-CoV-1, and Epstein-Barr Virus (EBV), a ubiquitous human herpesvirus that has been implicated in LC pathogenesis^14–16^ (complete list of genes included in the panel in **Table S1**); and (3) real-time PCR targeting the SARS-CoV-2 S1, S2, and N regions^17^.

Results from this suite of assays are summarized in **Fig 1** and described in detail below. A total of 12/44 (27%) of the LC group and 1/13 (7.7%) of the recovered group had evidence of SARS-CoV-2 persistence by any of the above bulk or in-situ tissue methods (p=0.26, Fisher exact test).

**Fig 1.**
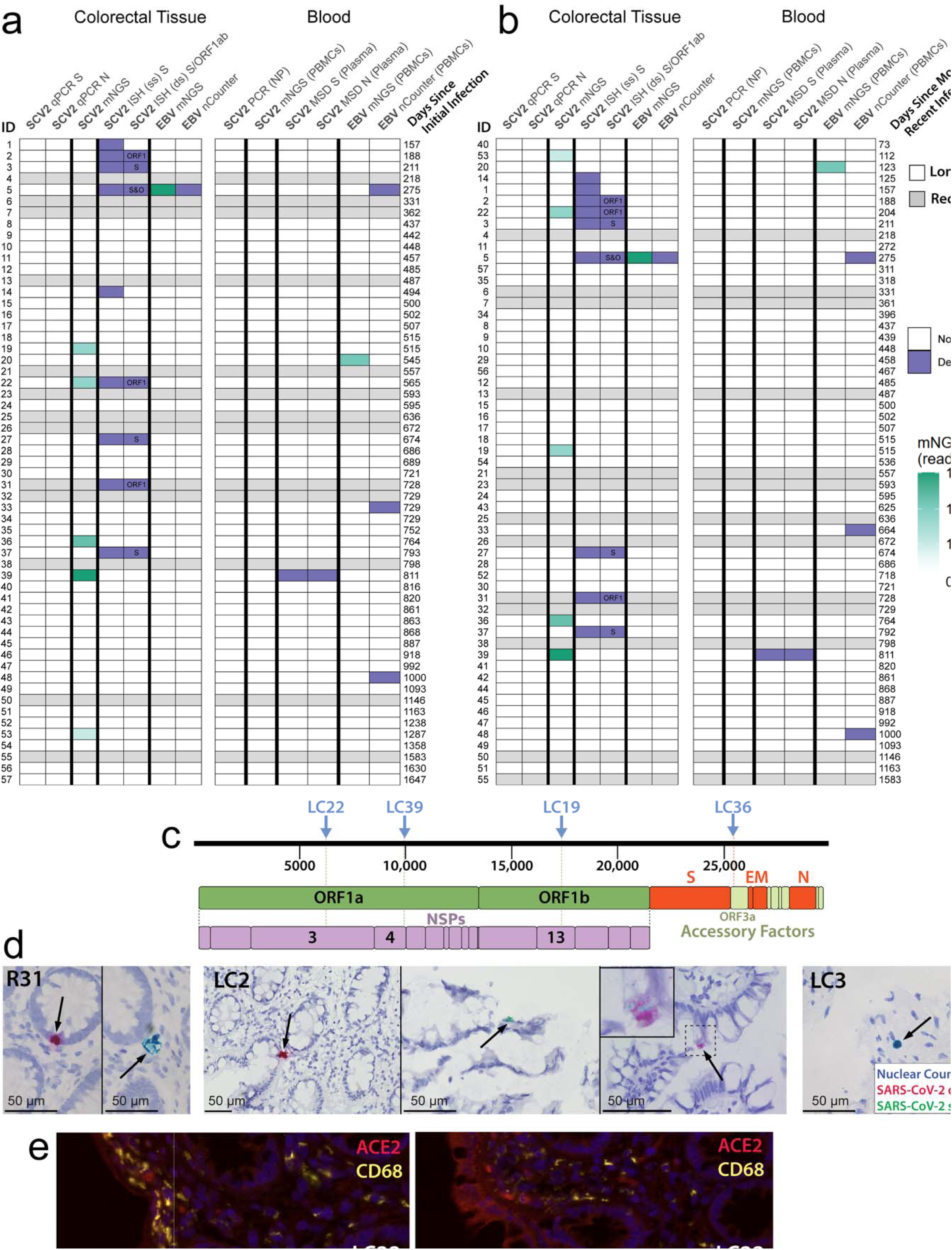
SARS-CoV-2 and Epstein-Barr Virus (EBV) detection in colorectal tissue and matched blood samples from participants with Long COVID (LC) and recovered (R) controls. A summary of viral detection sorted by time from initial infection (**a**) and most recent known or reported infection (**b**) to colorectal biopsy are shown. These graphs summarize results from tissue quantitative PCR (qPCR) for Spike (S) and Nucleocapsid (N) RNA, in-situ hybridization (RNAscope) of single-stranded (ss) and double-stranded (ds) viral RNAs targeting the S and ORF1a/b genomic regions, metagenomic sequencing (mNGS) with and without viral capture probe enrichment and nCounter, and circulating plasma S and N proteins. Panel (**c**) shows the sequence alignment locations of viral reads across the entire SARS-CoV-2 genome from each participant with a positive mGNS test. Panel (**d**) shows representative single-stranded (green) and double-stranded (red) SARS-CoV-2 RNA in colorectal sections from LC and recovered participants. Panel (**e**) shows example of duplexed ACE2 (red) and CD68 (yellow) fluorescent tissue immunohistochemical staining from participant LC33. CD68+ immune cells were observed scattered throughout the lamina propria (LP) in addition to residing in the sub-epithelium. ACE2 expression was low overall but was associated with epithelium and in subepithelial cells. PBMC = peripheral blood mononuclear cells, ORF open reading frame, NSP = non-structural proteins.

### In-situ detection of SARS-CoV-2

For the first five colorectal samples collected in our initial pilot study, we used two monoplexed fluorescent panels to identify single- and double-stranded SARS-CoV-2 Spike RNA, as previously reported^12^. Because tissue autofluorescence complicated interpretation, we transitioned to a duplex chromogenic method that simultaneously detected single-stranded Spike and double-stranded ORF1A/B RNA. We applied this assay to all 57 colorectal samples from participants with LC or recovered controls, including the five previously reported pilot samples. Two additional SARS-CoV-2 uninfected, pre-COVID-19 samples were used as negative controls and to establish non-specific background signal in each assay. This approach enabled colocalization of single- and double-stranded viral RNA without autofluorescence or spectral overlap. Regardless of the method used, multiple sections were assayed from each colorectal sample.

We identified single-stranded Spike RNA in colorectal tissue from 8 participants with LC (18.2%) and one recovered control (**Fig 1**). Seven of these nine samples, including the recovered sample, also contained detectable double-stranded ORF1A/B or Spike RNA. In the chromogenic assay, the frequency of positive ISH events was low, ranging from 28 to 245 counts per million nuclei, with a median of 59 counts per million.

We did not observe clear SARS-CoV-2 Spike protein IHC staining in any FFPE sample, although this approach has known sensitivity and specificity issues^18^.

### Metagenomic next-generation sequencing detection

We next analyzed the colorectal tissues using highly sensitive, unbiased mNGS approach for virus detection^19^, both with and without viral capture probe enrichment. SARS-CoV-2 RNA was detected by at least one of these approaches in tissue from five participants with LC (11.4%) but no recovered controls (**Fig 1a,b**). SARS-CoV-2 sequence counts and alignments are shown in **Fig 1c**. The detected viral sequence fragments mapped to multiple regions spanning the SARS-CoV-2 genome. Additional attempts to perform targeted genomic sequencing of RNA-positive tissues were unsuccessful because PCR amplification failed.

### Targeted viral RNA detection assays

We next applied our custom nCounter assay for human immune response profiling. After establishing conservative positivity thresholds based on background colorectal RNA signal in pre-pandemic samples and on SARS-CoV-1 transcript counts, no definitive SARS-CoV-2 RNA signal was detected. Importantly, we also found that RNA oversaturation of the assay produced false-positive signal. We therefore carefully controlled input concentrations to avoid assay oversaturation. EBV EBER1 and EBER2 transcripts, which are highly abundant in latently infected cells, were detected in one colorectal sample from a participant with LC (**Fig 1a**). qPCR assays in rectosigmoid tissues were negative in all participants.

### Spatial distribution of SARS-CoV-2 RNA in tissue

We observed two histological regions of SARS-CoV-2 RNA persistence in colorectal tissues. First, single-stranded Spike RNA and double-stranded ORF1A/B RNA were detected within the outermost epithelial layer (**Fig 1d**). Second, single-stranded RNA was detected in the sub-epithelium and lamina propria, whereas double-stranded RNA was not observed in these regions.

In our prior work, SARS-CoV-2 RNA in the lamina propria was associated with CD68-expressing myeloid immune cells^12^. However, in the current study, viral RNA was not consistently detected on contiguous sections used for immunophenotyping, precluding definitive assignment of the observed lamina propria signal to a specific cell type. We performed duplex ISH staining, however, on separate gut FFPE sections for ACE2 and CD68 proteins. Overall, ACE2 expression was low, as would be expected in distal colon/rectum compared to small bowel^20^, but localized to epithelium as well as non-epithelial cells in the sub-epithelial and other lamina propria tissue regions (likely fibroblasts). We also observed two distinct patterns of CD68 (myeloid/macrophage staining) including sub-epithelial and lamina propria localization. (**Fig 1e**).

### SARS-CoV-2 detection in peripheral blood

Multiple prior studies have attempted to identify circulating SARS-CoV-2 in plasma or cells in the peripheral blood^21–30^. We detected spike antigen in plasma from one participant with LC using the Meso Scale Discovery platform (MSD, Rockville, MD). Notably, SARS-CoV-2 RNA was also detected by mNGS in the paired colorectal tissue sample from this participant (**Fig 1a,b**).

However, SARS-CoV-2 RNA was not detected by nCounter or mNGS in peripheral blood mononuclear cells (PBMCs) from this or any participant. The limited concordance across tissue- and blood-based assays may reflect differences in the biological material measured by each assay, as well as the spatial heterogeneity of and/or low abundance of viral material.

#### Clinical correlates of SARS-CoV-2 detection

The median age and sex distribution of those with versus without SARS-CoV-2 detection were similar (43 vs 45 years, 20% of females and 27% of males (**Table S2**). The median time between the first known infection and colorectal sampling in those with SARS-CoV-2 detection by any method was 565 days (IQR 275-764), versus 725 days (505-903) in those without viral detection. The median time between most recent known infection and colorectal sampling in those with SARS-CoV-2 detection by any method was 275 days (IQR 188-728), versus 575 days (444-809) in those without viral detection. No clear pattern emerged in the clinical factors related to quality of life, LC symptom count, or specific LC symptoms. To further explore factors associated with detection of SARS-CoV-2 RNA in colorectal tissue, we performed univariate and multivariate logistic regression focused on the presence of LC and the following factors: (1) time since most recent SARS-CoV-2 infection, (2) sex, and (3) age (years). In univariate models, we identified a significant association between time since most recent infection and viral detection (p=0.045) but no other variable (all other p>0.05). Adjusted analyses including all factors in the model weakened the association between time since last infection and viral detection (adjusted p=0.077).

##### Analysis of the host immune response in blood and colorectal tissue

To assess the host immune response in LC compared to recovered participants, we performed differential expression analyses to a complementary set of sequencing-based, proteomic, and spatial methods to both colorectal tissue and paired PBMCs and plasma. In addition to the targeted transcriptomic panel described above (Ncounter), we employed (1) unbiased transcriptomic sequencing of colorectal tissue; (2) proteomic profiling of plasma samples using the Olink Reveal platform; (3) combined 5’ single-cell (sc)RNAseq with T-Cell Receptor (TCR)seq on a subset of 13 colorectal tissue samples; (4) high-dimensional 27-plex spectral flow cytometry on cryopreserved colorectal tissue; and (5) single cell spatial targeted transcriptomics (Xenium) on a subset of 6 FFPE colorectal tissue sections.

As detailed below, results from bulk tissue demonstrated broad and heterogeneous transcriptional dysregulation in colorectal tissue in the LC group, characterized by persistent inflammatory, proliferative, and metabolic remodeling. Single-cell analyses revealed specific functional reprogramming in colorectal tissue, particularly within epithelial, macrophage, fibroblast, and plasma cell populations. However, minimal changes in cellular composition were observed and we did not detect SARS-CoV-2 reads by single-cell RNA sequencing. Spatial analyses confirmed cell type-specific upregulation of immune pathways in macrophage and epithelial cells, although heterogeneity in inflammatory and immune responses was observed among the representative LC samples.

### Targeted transcriptome analysis

We characterized host transcriptional responses in RNA isolated from stabilized colorectal biopsy tissue and paired PBMCs, comparing participants with LC and recovered controls using two complementary approaches. First, we applied our custom nCounter assay. **Fig 2a,b** shows volcano plots, heatmaps of significant differentially expressed genes (DEGs) after false discovery rate (FDR) adjustments, and gene set enrichment analyses (GSEA) significance scores. Overall, we observed 240 significant DEGs in colorectal tissues when comparing participants with LC to recovered controls (complete list of DEGs in **Table S3**), with tight clustering of LC and recovered samples by gene expression patterns. One major recovered cluster and two major LC clusters were visualized. Five nanoString-defined gene pathways had robust Global Significance Scores (GSS^31^; GSS of >3) containing 7 to 54 individual significant genes in each set were identified (**Fig 2c**). Interestingly, all the top GSS pathways were downregulated overall (as defined by the Directed Global Significance Score; DGSS) and included major pathways associated with MHC class I antigen processing, mononuclear cell migration, lysosome activity, and phagocytosis. CCL8 and CCL13 were the most strongly downregulated individual genes, but others such as CD16a/b (NK/nonclassical monocyte markers), CCL4/eotaxin (eosinophil recruitment and bowel inflammation), CCL7, TLR3 (mediates detection of viral double-stranded RNAs), granzyme A (cytotoxicity) were also markedly downregulated. In contrast, IL1RN and S100A12, which are associated with immune activation and bowel inflammation, were the most highly upregulated genes.

**Fig 2.**
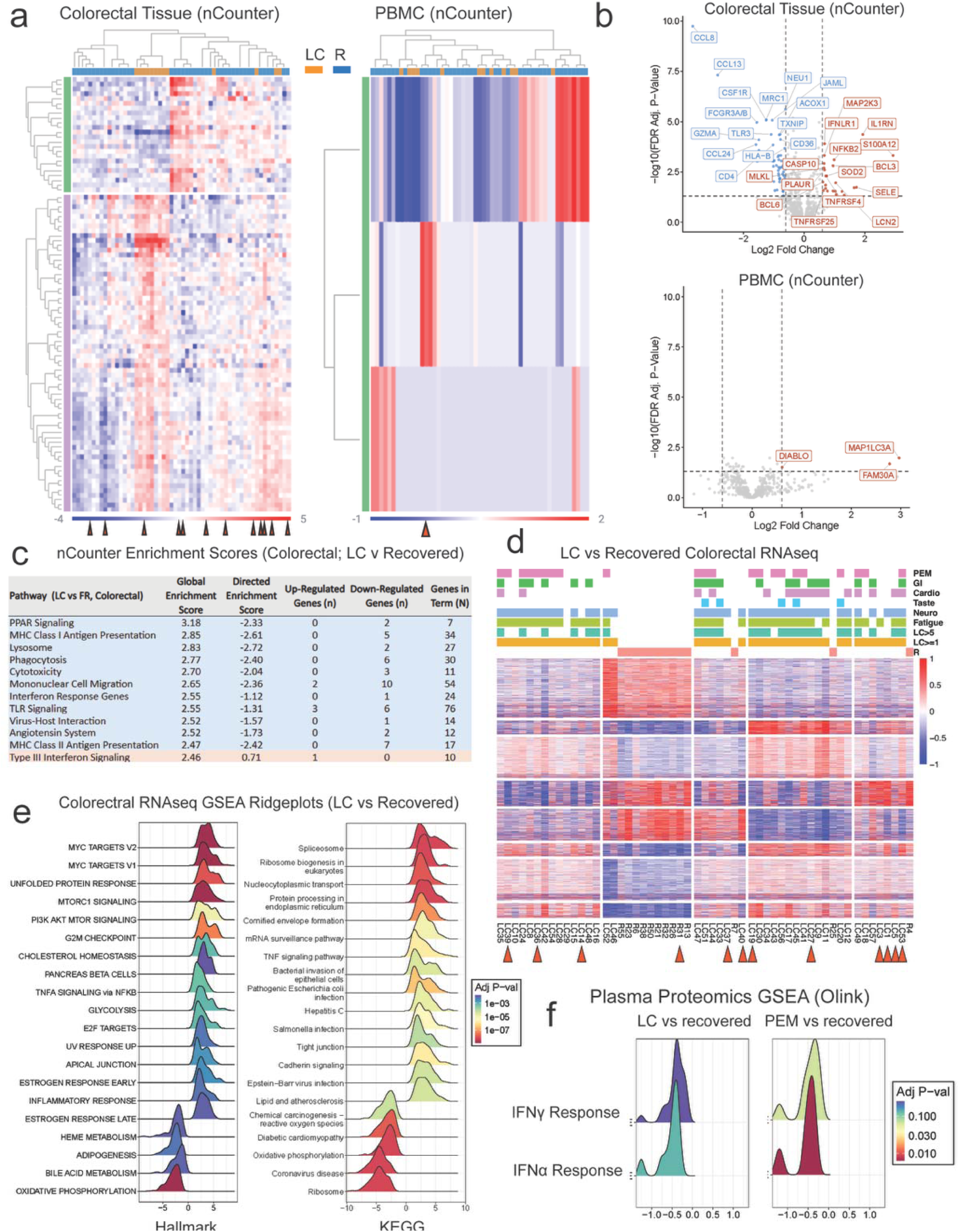
Targeted and bulk transcriptomic analyses of colorectal and peripheral blood mononuclear cell (PBMC) RNA and deep plasma proteomic profiling of Long COVID (LC) cases compared with recovered controls. Heat maps of significant differentially expressed genes using the nCounter targeted panel comprised of 807 gene transcripts representing viral immune responses and inflammation in colorectal tissue and PBMC are shown in (**a**). A list of all gene names included in the panel is shown in **Table S1**. 240 significant genes (DEG) were observed in tissue in two unique modules of genes (all significant gene names are shown in **Table S2**). Red arrows denote samples with positive SARS-CoV-2 detection by any method as in Fig 1. Volcano plots with top DEGs labeled (all significant after false discovery rate [FDR] adjustment) for colorectal tissue and PBMC are shown in (**b**). Global and Directed Enrichment Scores (GES, DES) from nCounter-defined transcriptomic pathways are shown in (**c**). A heat map of all 4757 FDR-adjusted DEGs from colorectal bulk tissue whole-transcriptome polyA-RNAseq in people with LC and recovered controls (n=55) is shown in (**d**). The top horizontal bars represent which participants experienced each LC symptom phenotype at the time of tissue sampling (PEM = post exertional malaise; GI = gastrointestinal, taste = smell and taste disturbances; fatigue; neuro = neurocognitive symptoms; LC>5 = participants experiencing greater than 5 LC symptoms, and LC>1 participants experiencing one or more LC symptom). 8 distinct gene modules were identified by non-hierarchical K-means clustering (row groups separated by white lines). A full list of significant DEGs is listed in Supplementary **Data File S1** and a heatmap with only the top 50 DEGs including gene names is shown in **Fig S2**. Red arrows denote samples in which SARS-CoV-2 was detected by any method. Panel (**e**) shows ridgeplots of gene-set enrichment analyses (GSEA) pathways (KEGG and Hallmark) with adjusted P values and ranking metrics on the x-axis defined by the signed log_10_ p-value. Results from deep plasma proteomic (Olink Reveal) analyses in people with LC versus recovered controls are shown in (**f**). Ridgeplots of protein enrichment analysis are shown for LC≥1 versus recovered and PEM versus recovered controls are shown. The interferon-α (IFNα) pathway was significantly downregulated in PEM vs recovered but no significance was observed when comparing the overall LC cohort or other individual LC phenotypes versus the recovered cohort.

Overall, the DEG and GSE analyses all point towards a decrease in gene transcriptional activity that is required for the immune system to recognize and eliminate virus-infected cells, although some inflammatory cytokines and NFkB activity were overexpressed in LC.

Importantly, the same gene expression analysis in matched PBMCs yielded only 3 significant DEGs, all of which were upregulated. These DEGs were not observed among those seen in the gut; **Fig 2b**, with no distinct clustering by disease phenotype. Furthermore, GSE analyses resulted in no robust pathways (GSS scores all <2.03). This suggests that biological differences using these analytic approaches may be more easily detectable in the tissues than in the peripheral blood.

### Unbiased transcriptomic analysis of colorectal tissue

Building on the targeted transcriptomic analyses using nCounter, we performed unbiased transcriptomic sequencing of colorectal biopsy samples to more comprehensively define the nature and extent of host gene responses. We identified 4,757 DEGs between participants with LC and recovered controls (n=55 as two LC participants did not have data available; **Fig 2d, Fig S1, Fig S2**), with clear separation and clustering between groups. Unsupervised hierarchical analysis further identified four distinct transcriptional clusters within the LC group defined by distinct patterns of DEGs as determined by K-means clustering (**Fig 2d**). These transcriptional clusters were not clearly associated with clinical phenotypes, including the presence of gastrointestinal symptoms (**Fig 2d**).

To further understand the unbiased RNAseq data, we again performed GSEA. We generated directional ridge plots to identify the top gene pathways and whether the genes in each pathway, taken together, were up or downregulated when comparing LC and recovered groups (**Fig 2e**, complete list of significant pathways in **Table S4**). Top significant upregulated Hallmark pathways in those with LC included MYC targets, MTORC1 and PI3K-AKT-mTOR signaling, TNFα signaling, glycolysis, inflammatory response, complement and several pathways involving cell cycling, metabolism and protein/hormonal responses.

Significantly downregulated Hallmark pathways in those with LC included heme metabolism, adipogenesis, bile acid metabolism, and oxidative phosphorylation. Applying Kyoto Encyclopedia of Genes and Genomes (KEGG) to the GSEA to better understand various disease processes associated with the DEGs, we observed increased pathways including mRNA surveillance, TNF signaling, and EBV infection (among others as shown in **Fig 2e**). Interestingly, the acute COVID-19 pathway was significantly downregulated in the LC versus recovered groups.

To further explore the relationship between SARS-CoV-2 detection and these immunologic changes, an additional differential expression analysis was performed comparing those with detectable virus (n=12) vs those without detectable virus (n=32) within the LC group. There were no significantly differentially expressed genes or enriched pathways, implying that changes in host immune response associated with LC occur independently of viral detection. A further comparison of LC participants who were hospitalized (n=3) vs those who were not hospitalized (n=41) during acute infection also revealed no significant findings.

Together, these data demonstrate broad and heterogeneous transcriptional dysregulation in colorectal tissue in Long COVID, characterized by persistent inflammatory, proliferative, and metabolic remodeling.

### Plasma proteomics

We performed deep proteomic profiling of immune activation and inflammation using the Olink Reveal platform^32^, which simultaneously tests 1,034 plasma proteins, in LC and recovered controls. We did not observe significant differences in protein expression between the overall LC and recovered cohorts; when examining specific phenotypes, we observed only one individual protein (Cathepsin H) upregulated in people with LC experiencing ongoing smell/taste disturbance compared to recovered individuals (**Fig S3**). Interestingly, we observed downregulation of IFNα and IFNγ Hallmark signaling pathways in the subset of people with LC experiencing PEM compared to recovered controls (**Fig 2f**).

### Colorectal single-cell RNA transcriptomics

To better understand the contribution of individual cell types within the rectosigmoid samples that may be leading to the bulk-transcriptome analyses, we performed combined 5’ single-cell (sc)RNAseq with T-Cell Receptor (TCR)seq on a subset of 13 participants including those with LC (n=9) and recovered controls (n=4) (**Table S5**). Among these 13 participants, we observed 12 major cell clusters as defined by Seurat including T cells, B cells, macrophages, three distinct plasma cell clusters, fibroblasts, endothelial cells, enteroendocrine cells, goblet cells, tuft cells, and colonic transit amplifying cells (TACs, which are rapidly dividing differentiating cells that are often in high abundance given rapid turnover of gut epithelium) (**Fig 3a**). Overall, there were no observed differences in the PCA cluster morphologies or recovered frequencies of cells within each cluster in LC versus recovered tissue groups (all p>0.05). Overall, there were only two significant DEGs within two distinct cell clusters that met individual FDR-adjusted statistical significance – ACE was significantly upregulated in intestinal goblet cells (1.7 Log_2_ fold change, Q=0.025) and Immunoglobulin J chain (JCHAIN) was significantly downregulated in macrophages **(Fig S4**). We did observe, however, strongly significant gene ontologies as defined by Hallmark and KEGG (**Fig 3b** shows all significant Hallmark pathways and the top 40 significant KEGG pathways; all KEGG pathways are shown in **Fig S5**).

**Fig 3.**
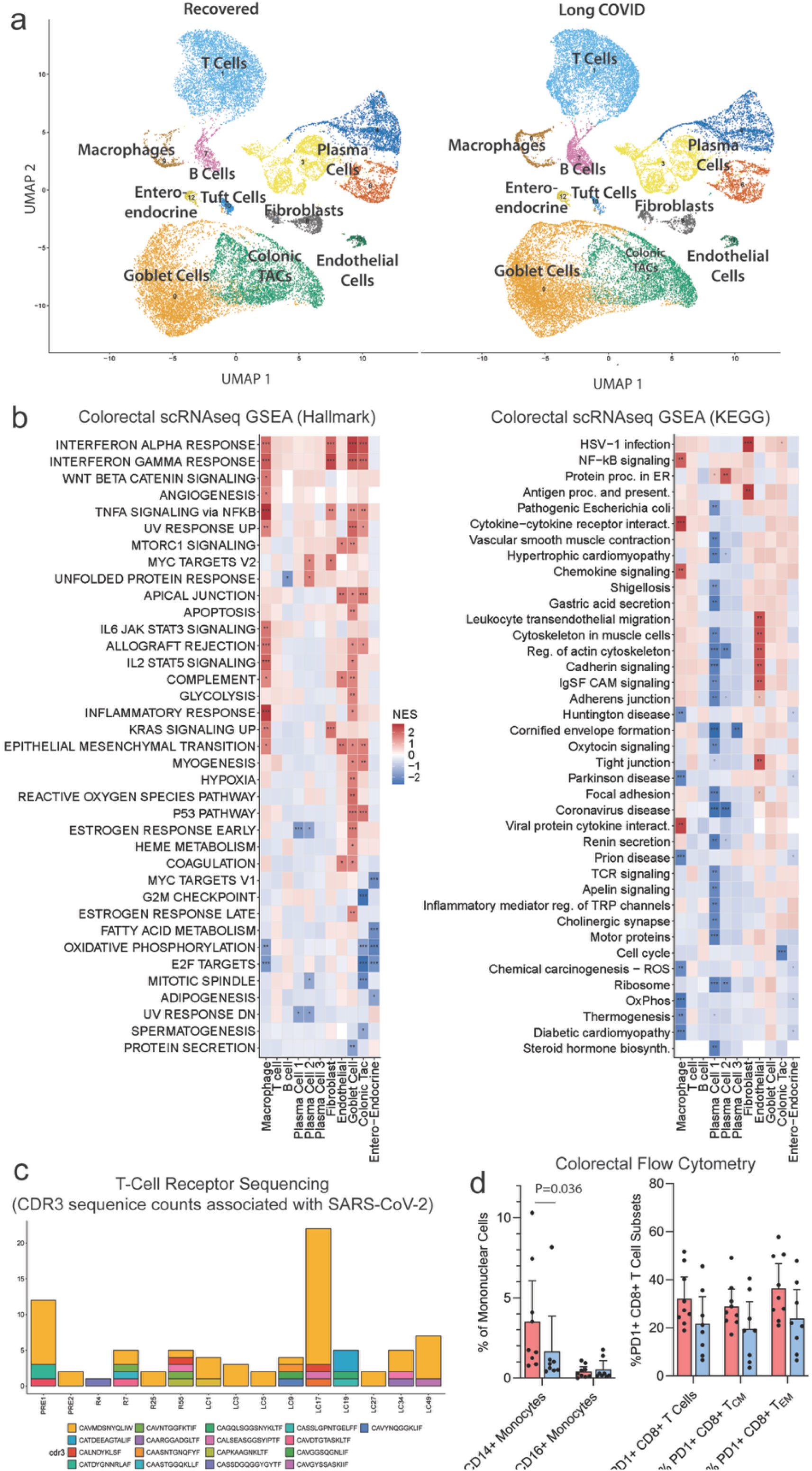
Unbiased single-cell transcriptomics, T-cell receptor sequencing, and spectral flow cytometric analyses of colorectal tissues in Long COVID participants and recovered controls. Uniform Manifold Approximation and Projections (UMAP) of unique cell clusters are shown in (**a**). Overall, there were no significant differences in the frequencies of each cluster-identified cell type or marked changes in overall UMAP morphologies between LC cases and recovered controls. Gene set enrichment analyses (GSEA; Hallmark and KEGG) of top significant gene expression pathways for each unique cell cluster are summarized in heatmaps are shown (**b**) shaded by Normalized Enrichment Scores (NES; upregulated in LC = red, downregulate in LC = blue). Significant different NES from FDR-adjusted analyses are labeled with asterisks (*<= 0.05, ** <= 0.01. *** <= 0.001, etc.). All significant Hallmark pathways are shown; KEGG pathways were filtered to only those with a p-value <= 0.01 (unfiltered version is shown in **Fig S4**). The percent of classical CD14+ monocytes and non-classical CD16+ monocytes and the percentage of PD1+ cells of various T cell subsets (total CD8+ T cells, central memory CD8+ T cells (Tcm) and Effector Memory CD8+ T cells (Tem) are show in (**c**). P values were calculated by non-parametric two-tailed, Mann-Whiteny U tests. Bars and lines represent mean and 95% confidence intervals. CDR3 T-cell receptor sequence counts with any confidence association with SARS-CoV-2 infection are shown (**d**) for T cell clusters across all participants included in the scRNAseq analysis. Two pre-COVID-19 samples were also included to understand the TCR landscape prior to infection (PRE).

We also identified upregulation of similar pathways in macrophages, which we previously demonstrated could harbor SARS-CoV-2 in the gut years after initial infection^12^, and to a lesser extent fibroblasts, which appear to express ACE2 in our IHC experiments (**Fig 1e**). Interestingly, B and T cells showed very little differences in any gene pathway and two of the three plasma cell clusters showed significant downregulation of pathways such as lymphocyte signaling pathways, coronavirus disease responses, and immunoglobin like cell adhesion. Of note, the absolute number of plasma cells and lymphocytes were higher than the number of myeloid immune cells and fibroblasts in colorectal tissue (**Fig 3a**), which may explain the bulk nCounter and RNAseq analyses showing downregulation of many adaptive and cytotoxic immune response genes as above.

In addition, we analyzed the single-cell RNAseq data using the clinically validated SURPI mNGS analysis pipeline for viral infection^33^ but identified no reads that aligned to SARS-CoV-2. SURPI applies stringent filtering and taxonomic classification criteria to distinguish true viral sequences from spurious alignments and other causes of false-positive viral read detection that can confound other viral sequence alignment algorithms^19,33–35^.

Together, these findings indicate that LC is associated with specific gene expression reprogramming in colorectal tissue, particularly within epithelial, macrophage, fibroblast, and plasma cell populations, despite minimal changes in cellular composition and no definitive detection of SARS-CoV-2 reads by single-cell RNA sequencing.

### Colorectal single-cell T cell receptor (TCR) sequencing

Complementarity-Determining Region 3 (CDR3) is the most hypervariable loop of the TCR and the primary determinant of T cell antigen specificity. CDR3 sequences were obtained from single-cell TCRseq data to determine TCR landscapes of SARS-CoV-2-associated sequences between LC and recovered groups. **Fig 3c** shows the frequencies of CDR3 sequences that have some association with SARS-CoV-2 infection, including those with some level of confidence that the CDR3 region is elicited by SARS-CoV-2 antigens. We observed one LC participant who had a high number of one CDR3 sequence compared to the remaining participants, but this sequence was also identified in the pre-COVID-19 controls, although to a lesser extent (**Fig 3c**).

### Flow cytometric analysis of colorectal cells

To characterize immune-cell phenotypes and protein expression in colorectal tissue, we performed high-dimensional 27-plex spectral flow cytometry on collagenase-digested, thawed cryopreserved tissue from participants with LC and recovered controls, including cells isolated from the intraepithelial and laminal propria fractions. Samples were selected based on the availability of sufficient bulk cryopreserved tissue, while maximizing overlap with participants who were included in the scRNAseq experiments (**Table S6**). This panel was designed to phenotype γδT cells, CD8+ and CD4+ T cell memory subsets (naive, central memory, effector memory, terminally differentiated), T cell activation and immune checkpoint [PD-1] markers, IgA- an IgM-producing memory B/plasma cells, NK cells (immature cytokine producing, mature cytotoxic), classical and non-classical monocytes, and basophils (complete gating strategy outlined in **Fig S6-8**).

Participants with LC had a significantly higher percentage of CD14+ classical monocytes in colorectal cells than recovered controls (p=0.036). We also observed a non-significant trend towards a higher percentage of PD1-expressing CD8+ T cells in the LC group (P=0.076; **Fig 3d**). We did not observe significant differences in the frequencies of memory B/plasma cells expressing IgA or other cell types in the flow panel.

### Single-cell spatial profiling (Xenium) of colorectal tissues

To visualize gene expression data across space, we developed a custom Xenium (10X) panel to perform spatial RNA profiling in FFPE sections of colorectal tissues from 4 LC and 2 recovered controls (participant demographics are found in **Table S7**, all transcripts in the panel are found in Table **S8**). The four LC samples (2 males, 2 females) were chosen based on the presence of detectable SARS-CoV-2 by any method and the availability of sufficient tissue for additional analyses. One male and one female recovered control were also selected as comparators. We modified a human pan-tissue off-the-shelf panel to include custom human and SARS-CoV-2, EBV, and CMV transcripts across multiple viral genomic regions. Unique single cell clusters were defined based on transcripts detected within each individual cell across the entire tissue. We mapped the tissue distribution of gene expression pathways identified from the scRNAseq analyses with single cell resolution.

Specifically, Pathway Activity Scores (PAS) were calculated for each individual cell and visualized spatially in 2-dimensions grouped by each unique cell cluster. **Fig 4** shows a representation of these pathway activity scores in macrophages, epithelial cells, and fibroblasts. We observed substantial biological heterogeneity within the LC group, but two participants had clearly different intensity of PAS differential regulation compared to recovered controls. Interestingly, areas of upregulated and downregulated pathway activity were broadly distributed across whole tissue sections and did not show focal localization to areas around sites of detectable viral RNA.

**Fig 4.**
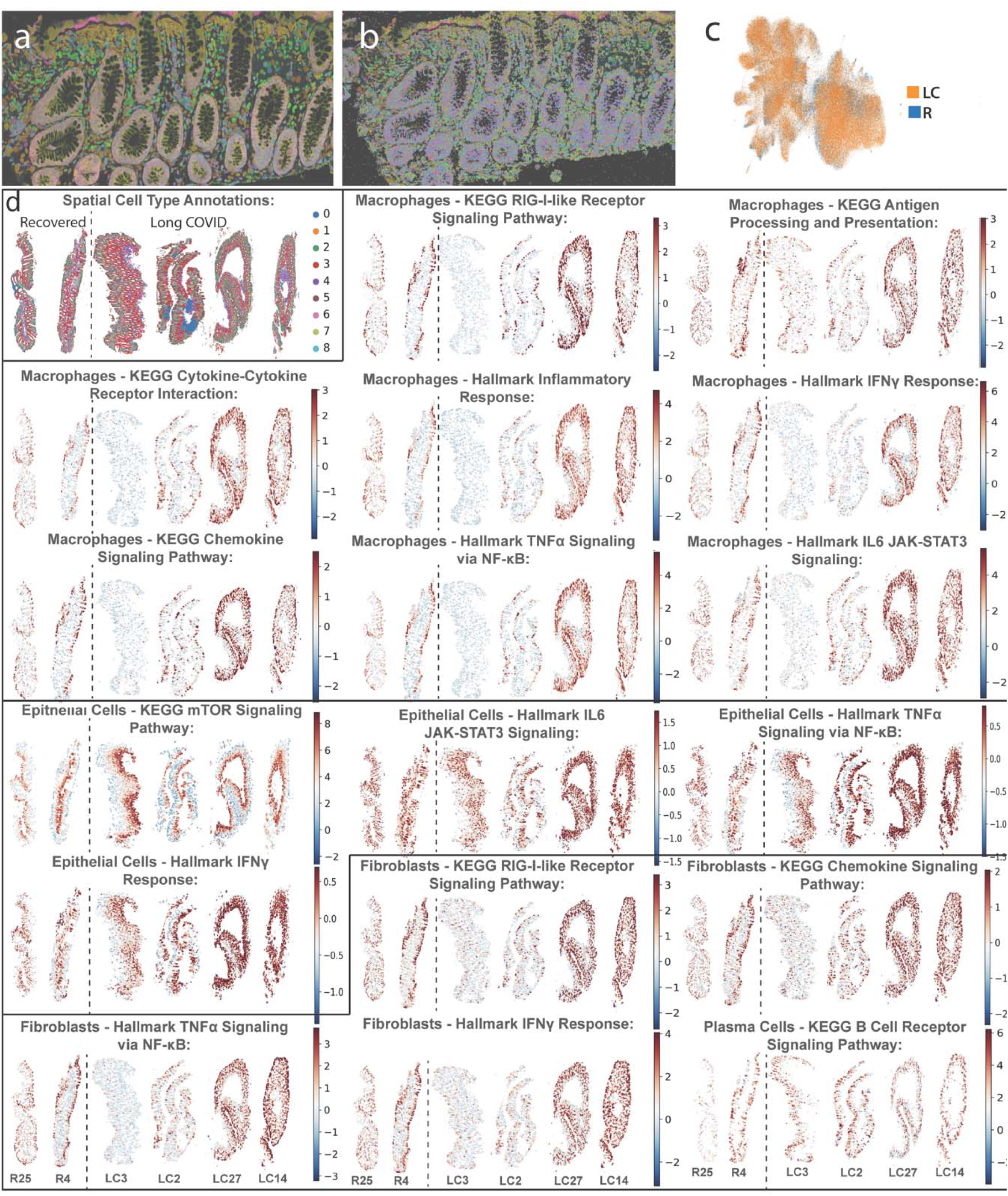
Spatial mapping of gene pathway activity from spatial transcriptomic analyses. The Xenium (10X) platform was used to visualize significant gene pathways of interest as identified in bulk and single-cell RNA sequencing experiments across a sub-set of Long COVID (LC; n=4) and recovered (R; n=2) participants. A unique cell type (cluster) is assigned to each individual cell across space using a combination of immunohistochemical staining for nuclei, cell wall material and intracellular RNA (**a**) as well as individual transcripts shown as different colored pixels mapped to within each cell (representative example shown in **b**). A UMAP of single cells, colored by condition. **c**). Cell type annotations across all cells in each tissue are shown (**d**) from the 4 LC and 2 recovered participants. Cell clusters included: 0 = T cells, 1 = Mast cells, 2= Macrophages, 3 = Epithelial cells, 4 = Smooth Muscle Cells, 5 = Colonocytes, 6 = Fibroblasts, 7= Plasma Cells, 8 = Endothelial Cells. Gene Activity Scores (GAS) were calculated for each individual cell using the Xenium transcript data for significant gene pathways identified from the single-cell RNAseq experiments (Fig 3). Representative GAS scores of each cell for macrophages, epithelial cells and fibroblasts are also shown across 2-dimensional space.

## DISCUSSION

In this study, we performed high-dimensional profiling of colorectal tissue and peripheral blood from a deeply characterized cohort of 44 people with Long COVID and 13 recovered controls. Using complementary sequencing-based, proteomic, and spatial detection methods, we observed that SARS-CoV-2 persistence is enriched in LC compared to recovered participants, with higher prevalence in tissue than in blood. Host immune response analyses from bulk tissue showed a marked decrease in gene transcriptional activity required for viral clearance. Single-cell analyses revealed robust upregulation of innate inflammatory pathways in macrophages, epithelial cells, and, to a lesser extent, fibroblasts, contrasting with a downregulation of lymphocyte signaling pathways and viral responses in plasma cells. Finally, spatial transcriptomic analyses demonstrated that upregulation of inflammatory signaling in macrophages, epithelial cells, and fibroblasts was distributed throughout the colorectal tissue, rather than localized to the site of viral persistence. Overall, these findings suggest that Long COVID is defined by a tissue-based transcriptional environment in which SARS-CoV-2 activates innate myeloid immune signaling, driving chronic inflammation while leading to dysregulation of pathways responsible for immune-mediated clearance of infected cells.

A key advance of this study is the coexistence of sustained innate immune activation with attenuation of the adaptive immune pathways required for effective elimination of virus-containing cells. Across transcriptional and spatial analyses, macrophages, other myeloid populations, and fibroblasts exhibited programs consistent with ongoing viral sensing and inflammatory signaling, including activation of TLR-, RIG-I- and TNF-associated pathways. At the same time, pathways involved in phagocytosis, antigen processing and presentation, and cytotoxic lymphocyte function were broadly downregulated in bulk tissue analyses. These abnormalities occurred with relatively limited changes in overall tissue cellular composition, indicating an altered cellular state rather than broad changes in immune cell abundance. The resulting picture suggests an uncoordinated immune response in which inflammatory sensing remains active while the mechanisms needed to identify and eliminate the source of that stimulation are impaired.

Although immune dysregulation represents the most consistent finding of this study, the relationship between these abnormalities and viral persistence remains unresolved. We detected SARS-CoV-2 RNA in colorectal tissue from 12 of 44 participants with LC and 1 of 13 recovered controls, indicating that detectable viral persistence is enriched in LC but is neither universal nor entirely specific to the condition. Importantly, the prevalence of viral persistence is likely underestimated in this study, given that (1) the extremely low abundance of viral RNA relative to host transcripts makes detection technically challenging, and (2) endoscopic colorectal biopsies sample only a minute fraction of the extensive gastrointestinal tract. Viral reservoirs may be distributed focally, intermittently, or preferentially in anatomical sites that were not sampled. More comprehensive sampling across different regions of the gastrointestinal tract will be required to define the true distribution of persistent viral material.

The histological localization of the detected viral RNA and host immune responses provide additional biological context. Single- and double-stranded viral RNA were detected within colorectal epithelial cells, which undergo rapid turnover on the order of several days^36^. In the absence of a new infection, this observation is suggestive of ongoing viral replication or re-seeding of epithelial cells. We also detected double-stranded viral RNA within the lamina propria, which we have previously demonstrated to be localized within CD68+ myeloid immune cells^12^. In contrast, we were unable to reliably assess viral protein because currently available immunohistochemical assays lacked sufficient specificity for inclusion in this study. Overall, these findings suggest ongoing active viral activity in colorectal tissues months to years after most recent known infection, although we cannot exclude the possibility of more recent, subclinical infection leading to some of these observations.

Regardless of the detection of SARS-CoV-2 RNA, several explanations could account for the striking features of tissue-based immunologic pathways observed in LC. One possibility is that persistent virus, even if not detected on biopsy, directly drives chronic innate immune activation. This hypothesis is consistent with the observed enrichment of viral sensing pathways in specific cell types (macrophages, epithelial cells, and fibroblasts). Our spatial transcriptomic analyses demonstrate that the host viral responses identified in LC extend well beyond discrete sites of viral persistence, encompassing widespread alterations across the intestinal tissue microenvironment. An alternative theory is that other pathophysiologic processes, perhaps related to the severity of the initial infection, may lead to a higher tissue viral burden, which initiates long term immune dysfunction independent of viral persistence. Between these two models, our observation that innate immune activation is sustained months to years after the most recent infection implies the ongoing presence of immune stimuli, i.e. persistent virus, even if it cannot be readily detected. However, we also note that despite anecdotes suggesting improvement in LC with antiviral therapy^37–41^, randomized trials employing small molecule antivirals or monoclonal antibodies to date have not demonstrated a clear benefit among those with broadly defined LC^42–44^.

The specialized immune environment of the gastrointestinal tract may help explain how viral persistence can exist alongside ongoing inflammation and ineffective clearance. The gastrointestinal tract is a uniquely tolerogenic immune environment that restrains adaptive immune activation while maintaining barrier homeostasis through IgA-producing plasma cells and other regulatory populations^45^. Our findings raise the possibility that SARS-CoV-2 exploits, or is inadvertently protected by, this tolerogenic environment. SARS-CoV-2 has been shown to use syncytia formation during acute infection to evade interferon- and antibody-mediated antiviral activity in vitro and in vivo,^46^ raising the possibility that persistent infection may be further facilitated by cell-to-cell spread through syncytia, which could reduce immune recognition while continuing to trigger local innate inflammatory signals. Plasma-cell-associated programs and other local signals may limit the recruitment, activation, or effector function of cytotoxic T cells and natural killer cells, allowing virus-containing cells to persist despite continued innate immune sensing. Spatial analyses further suggest that macrophages and fibroblasts may act as important organizers of this inflammatory niche, coordinating persistent innate activation within the surrounding tissue. This interpretation remains hypothetical, as the present data cannot establish causation. Nevertheless, the co-occurrence of regulatory immune programs with impaired antigen presentation and cytotoxic responses is a plausible tissue context in which mucosal immune homeostasis may come at the cost of inability to achieve full viral clearance.

Although recent blood-based studies have identified persistent antigenemia and immune dysregulation in LC^15,21–30^, our findings demonstrate that these biological processes are considerably more pronounced within the colorectal tissue microenvironment than in the blood. These observations suggest that key mechanisms of LC may be anatomically compartmentalized and only weakly reflected in the circulation. This has important implications for studies that rely exclusively on blood-based measurements, which may underestimate persistent viral activity or fail to capture the immune processes operating within affected tissues. The lack of concordance between tissue and blood findings here underscores the urgency of efforts to relate blood- and tissue- based findings and develop non-invasive, scalable, methodologies for LC diagnostics, which are now underway.

At the same time, it remains unclear how localized gastrointestinal persistence and immune dysregulation would produce the multisystem manifestations of LC. Potential mechanisms by which this could occur include generation of circulating inflammatory mediators, microbial or viral products released through impaired barrier function^47,48^, or neural signaling through vagal afferents^49,50^. Alternatively, gastrointestinal persistence may represent one component of a broader, anatomically distributed reservoir rather than the primary source of systemic disease. Determining whether the gut directly drives distant organ dysfunction, contributes indirectly through systemic immune or neural signaling, or serves as a marker of persistence elsewhere will require longitudinal studies linking tissue findings to circulating mediators, barrier integrity, autonomic signaling and organ-specific outcomes.

We did not identify robust associations between biological endotypes in the colorectal tissue and LC clinical phenotypes. Although the modest sample size limited power to detect such relationships, this disconnect has important implications for disease classification and clinical trial design. A shared tissue process may produce heterogeneous manifestations depending on host susceptibility, involvement of other organs, and downstream inflammatory pathways. Conversely, similar symptoms may arise from distinct biological mechanisms. The clinical phenotype may therefore be an unreliable surrogate to predict who might respond to a mechanism-specific intervention. Whether symptom-defined subgroups necessarily represent distinct biological entities remains an open question. As LC trials increasingly enroll or stratify participants according to pre-defined symptom clusters, our findings support also the consideration of approaches that enroll or stratify participants based on biomarkers of viral persistence, tissue inflammation, or immune dysfunction.

Although single-therapeutic trials targeting viral persistence have to date been negative^42–44^, our finding of innate immune activation and impaired cytotoxic clearance in colorectal tissue regardless of the presence of viral detection provides a rationale for interventions that address both the reservoir and the tissue immune environment that allows it to persist. Antiviral therapy may suppress replication or reduce the production of immunostimulatory viral material, but could be insufficient if there exists a long-lived reservoir of virus-containing cells that are not maintained by active replication^51^. Conversely, nonspecific immune stimulation could intensify inflammation without restoring coordinated antiviral activity^4^. Rational treatment strategies may therefore require approaches that focus on boosting infected cell recognition and clearance, perhaps via immunotherapeutic strategies that seek to improve antigen presentation, cytotoxic T-cell activity, or natural killer-cell function. These findings also raise potential preventive implications. Earlier treatment of acute infection and prompt treatment or prevention of reinfection could reduce the establishment or replenishment of tissue reservoirs^52^, providing a mechanistic hypothesis for the observed benefit of trials of early antiviral treatment^53–55^ that could be tested in prospective trials to prevent post­acute consequences following SARS-CoV-2 infection.

Strengths of this study include its relatively large cohort for an invasive tissue-based investigation, deep clinical characterization, systematic assessment of intervening infections, and paired analysis of colorectal tissue and peripheral blood. The inclusion of carefully characterized individuals who had recovered after COVID-19 was particularly informative, revealing that gastrointestinal viral persistence is enriched in LC but is not entirely specific to symptomatic illness, thereby allowing a more nuanced assessment of its biological significance. Several limitations should be considered. This was a cross-sectional study, precluding assessment of the temporal relationships among viral persistence, tissue immune dysregulation, and clinical illness. The sample size remained modest for analyses relating tissue endotypes to clinical phenotypes, and much larger studies will be needed to test whether such relationships exist. Sampling was restricted to the colorectum, and persistence may involve other gastrointestinal regions or organs – a negative colorectal biopsy cannot establish the absence of viral persistence elsewhere in the GI tract or body. Detection of viral RNA or protein does not prove replication-competent virus, and low viral abundance limited the yields from amplification and genomic sequencing. Heterogeneity in the timing, number and variants of previous infections may also have contributed to biological variability. Finally, the observational design cannot establish that gastrointestinal persistence causes the associated immune abnormalities or systemic illness, and the transcriptomics results require functional validation.

In summary, this study supports a model of LC characterized by a distinct tissue immune environment with sustained innate immune activation and dysfunction consistent with ongoing viral persistence. Despite direct viral detection in only a subset of participants, we identified broadly distributed single-cell transcriptional responses across the tissue distinguishing LC from recovery. Our findings suggest the need for the field to shift toward understanding not just in whom persistence can be detected but also how persistence interacts with tissue immune regulation and how that interaction may contribute to chronic disease. The observation that tissue-based viral persistence also occurs in some recovered individuals further reinforces that persistence alone is unlikely to be solely responsible for LC, and that host immune responses may determine whether persistence is tolerated or progresses to a clinically apparent disease state. These findings provide rationale for ongoing efforts to assess antiviral strategies but also support therapeutic approaches that restore or redirect cytotoxic immune surveillance, or treatment combinations targeting both the persistent reservoir and tissue-based immune dysfunction. Ultimately, such interventional studies will be required to establish whether these processes are causally related to LC and whether targeting these mechanisms can prevent or treat the condition.

## FOOTNOTES

### Author Contributions

JDK, JNM, SGD, CYC, MS, MJP, and TJH conceived and designed the study. BL, DM, JV, TD, JC, CYC, MJP, and TJH developed the methodology and performed formal analyses. BL, JV, AER, AT, LG, EV, UV, BAP, TRF, YF, ML, DR, MD, BA, JL, AL, NR, CBS, HNM, CW, NK, RT, SP, MT, ZT, ZL, GS, AF, NS, VK, EAF, RH, JDK, JNM, SGD, CYC, MS, MJP, and TJH conducted the investigation. Data curation was performed by JV, AER, AT, LG, EV, UV, BAP, TRF, YF, ML, DR, MD, BA, JL, AL, NR, CBS, HNM, NK, VS, RT, SP, MT, ZT, GS, AF, NZ, NS, VK, EAF, RH, JNM, MJP, and TJH. Resources were provided by AER, YF, ML, DR, MD, JL, RH, JDK, JNM, SGD, CYC, MS, MJP, and TJH. BL, DM, JV, TD, JC, and TJH developed software and computational tools. Validation was performed by BL, JV, and TJH. Visualization was performed by BL, DM, JV, AT, TD, JC, CYC, MJP, and TJH. EAF and RH managed project administration. Funding was acquired by SGD, MJP, and TJH. JNM, SGD, CYC, MS, MJP, and TJH supervised the study. BL, DM, JV, AER, TD, JC, MJP, and TJH wrote the original draft of the manuscript. All authors reviewed, edited, and approved the final manuscript.

### Funding

PolyBio Research Foundation, NIH Common Fund U01AT012993, NINDS R01NS136197, NIAID R01AI93318. Research reported in this was also supported by the UCSF-Bay Area Center for AIDS Research, an NIH-funded program under award number P30AI027763 which is supported by the following NIH Institutes and Centers: NIAID, NCI, NIDA, NICHD, NHLBI, NIA, NIMHD, NINR, NIDDK, NIDCR.

### Disclosures

MJP has received consulting fees from Gilead Sciences, AstraZeneca, BioVie, Apellis Pharmaceuticals, BioNTech, and TechImmune, travel support from Invivyd, and research support from Aerium Therapeutics, Shionogi, ImmunityBio, and Enanta Pharmaceuticals, outside the submitted work. TJH receives research support from Shionogi and ImmunityBio and has consulted for ImmunityBio and Roche.

## Acknowledgements

We acknowledge the study participants, their caregivers, and their medical providers. We also thank Elnaz Eilkhani for regulatory support; Viva Tai for her general coordination of the research program; the UCSF LIINC and RECOVER teams for participant referral, recruitment, and study coordination; the San Francisco General Hospital Gastroenterology nurses for their assistance with colorectal tissue collection and participant care; and the UCSF Specimen Processing and Banking Subcore, including Salman Mahboob and Billy Huang, for overseeing biospecimen processing and storage. We further acknowledge the clinical and procedural teams involved in colorectal tissue collection, as well as the laboratory and core-facility personnel who supported tissue processing, imaging, sequencing, flow cytometry, and transcriptomic analyses. We are grateful to Amy Proal from PolyBio Research Foundation for her advice and input on the project. CYC is a co-founder of Delve Bio and on the scientific advisory board for Delve Bio and Romix Biosciences; he also receives research support from Delve Bio and Abbott Laboratories, Inc. and is an inventor on US patent 11380421, “Pathogen detection using next-generation sequencing”, under which algorithms for taxonomic classification, filtering and pathogen detection are used by SURPI+ software.

## METHODS

### 1. Participants

#### 1.1 LIINC Cohort

Participants were enrolled in the University of California, San Francisco (UCSF)-based Long-term Impact of Infection with Novel Coronavirus (LIINC) study (NCT04362150), an ongoing prospective cohort investigating the long-term clinical and biological consequences of SARS-CoV-2 infection. Adults with documented SARS-CoV-2 infection confirmed by antigen or nucleic acid testing were eligible for enrollment at least 14 days after symptom onset. Participants underwent standardized study visits approximately every four months that include interviewer-administered assessments of symptoms, quality of life, medical history, additional SARS-CoV-2 infections, vaccination history, and peripheral blood collection. Participants were asked to rate their overall health in the preceding 7 days on a 100-point visual analog scale (VAS), with 0 representing the worst and 100 the best health they could imagine. Procedures have been previously described in detail^56^.

Within LIINC, LC is defined as one or more new or worsened symptoms beginning after SARS-CoV-2 infection, persisting for at least 3 months, and not better explained by an alternative diagnosis. This definition is consistent with established LC case definitions, including those developed by the World Health Organization^13^ and the National Academies of Sciences, Engineering, and Medicine^57^. Recovered controls were LIINC participants who reported resolution of symptoms attributable to COVID-19 following their SARS-CoV-2 infection.

LIINC participants were screened for eligibility for the gastrointestinal tissue substudy. Screening included review of medical history and laboratory testing to exclude conditions associated with increased procedural risk, including clinically significant coagulopathy, thrombocytopenia (platelet count <100,000 µl⁻¹), neutropenia, current anticoagulant use, or inability to temporarily discontinue aspirin or non-steroidal anti-inflammatory drugs according to the study protocol.

In addition to participants in whom an initial analysis of colorectal biopsy specimens using RNAscope was reported^12^, several participants provided colorectal tissue in relation to their participation clinical trials within our research program^44^. In such cases, samples for this study were either obtained prior to baseline colorectal sampling in the clinical trial (n=17; i.e., a sample taken prior to pre- and post-intervention biopsies) or from baseline, pre-intervention timepoints (n=14); no post-intervention biopsies were included in this analysis. There was minimal scientific overlap with analyses performed in the clinical trial settings and the analyses reported in this manuscript.

#### 1.2 Descriptive statistics

Descriptive statistics are reported for the demographics of the cohort. Continuous variables were compared using the Wilcoxon rank-sum test. Categorical variables were compared using two-sided Pearson’s chi-square or Fisher’s exact tests, as appropriate. A p-value <0.05 was considered statistically significant.

Clinical correlates of detection were assessed by Binary logistic regression analyses performed using SPSS (vs 32, IBM) including the constant in the model.

#### 1.3 Colorectal biopsies

Following informed consent and screening procedures, participants self-administered two saline enemas approximately 2 h before the procedure. Participants of childbearing potential underwent urine pregnancy testing prior to tissue collection; those with a positive pregnancy test were excluded from the study procedure. Flexible sigmoidoscopy was performed, and random rectosigmoid mucosal biopsies were collected approximately 30 cm from the anal verge using 3.7mm disposable biopsy forceps. Collected tissue was placed in pre-labeled tubes containing either RPMI with 15% fetal bovine serum (R15) media for immediate transport to the laboratory for cryopreservation, nucleic acid stabilization or flash freezing for subsequent cellular analyses or freshly prepared paraformaldehyde for histological and spatial analyses.

Participants included in this analysis underwent flexible sigmoidoscopy between October 2020 and May 2025.

#### 1.4 Tissue Processing

Rectosigmoid biopsy samples collected in paraformaldehyde were fixed for approximately 24 h, dehydrated, and subsequently paraffin embedded. Samples collected in tissue culture medium were transported to the processing laboratory at 4 °C and processed within 1 h of collection. Tissue designated for cellular analyses was cryopreserved in fetal bovine serum containing 20% dimethyl sulfoxide and stored in liquid nitrogen according to previously described methods^15^. Processing procedures specific to individual downstream assays are described below. RNA was extracted from rectal tissue using the AllPrep DNA/RNA Mini Kit (QIAGEN® 80204) per manufacturer instructions.

#### 1.5 Blood and nasopharyngeal swab collection

On the day of sigmoidoscopy, participants underwent peripheral blood collection. Blood was processed as serum, plasma, and cryopreserved PBMCs. In most cases, participants provided a nasal swab for SARS-CoV-2 PCR testing to confirm the absence of nasopharyngeal shedding or occult re-infection at the time of gut biopsy.

#### 1.6 Ethics

The LIINC study and gastrointestinal tissue sub study were approved by the UCSF Institutional Review Board. All participants provided written informed consent for the parent LIINC study and the gastrointestinal tissue sub study.

### 2. RNAscope in-situ hybridization (ISH)

For the initial 5 participants with colorectal samples collected, a fluorescent ISH assay was used for in situ detection of viral RNA as previously described^12^. Given issues with autofluorescence and spectral overlap in this first experiment, we decided to implement a chromogenic ISH assay which was performed on samples from all study participants. The manual RNAscope® 2.5 HD Duplex assay (Advanced Cell Diagnostics, catalog no. 322430) was used to identify SARS-CoV-2 single-stranded spike RNA [probe-V-nCoV2019-S (catalog no. 848561)] and double-stranded orf1ab RNA [V-nCoV2019-orf1ab-sense (catalog no. 859151-C2)] in situ. Paraffin-embedded tissue blocks were sectioned at 5 μm and mounted onto SuperFrost Plus slides, which were stored in a desiccant chamber prior to staining. A minimum of three FFPE rectosigmoid tissue biopsies from each participant were used for RNAscope® experiments, along with comparative uninfected tissue, and were mounted on the same slide to control for batch effects from processing, staining, microscopy, and image analysis. Slides were baked in a dry air oven at 60°C for 1 hour, deparaffinized twice in 100% xylene (5 min), and dehydrated twice in 100% ethanol (1 min). To block endogenous peroxidase activity, slides were incubated with hydrogen peroxide for 10 minutes. Next, heat-induced epitope retrieval was performed for 18 minutes at 100°C using Target Retrieval Reagent (ACDBio). Protease digestion was accomplished by treatment with Protease Plus solution (ACDBio) for 30 min at 40°C. Hybridization was then performed with a duplex cocktail of probe-V-nCoV2019-S (catalog no. 848561, 1:1 dilution) and probe-V-nCoV2019-orf1ab-sense (catalog no. 859151-C2, 1:50 dilution) for 2 hours at 40°C. Following hybridization, amplifications one through ten, as well as detection of the red and green channels, were performed according to the manufacturer’s original protocol. Following detection of the green channel, slides were counterstained with Gill’s Hematoxylin I (StatLab, catalog no. HXGHE1LT, 1:2 dilution), blued with .02% ammonia water (Thermo Scientific Chemicals, catalog no. 1336-21-6), and mounted using VectaMount Permanent Mounting Medium (H-5600-60).

Images were captured using the Keyence BZ-X810. FFPE rectal tissues from pre-pandemic participants, as well as FFPE lung tissue from SARS-CoV-2 infected K18-hACE2 mice, were run as negative and positive controls, respectively. Sections from all FFPE blocks were also stained with a human housekeeping gene cocktail (POL2RA, PPIB) to assess block RNA integrity and assay performance.

### 3. Immunohistochemistry (IHC)

#### 3.1 Spike protein IHC

FFPE rectal biopsy specimens (n=36) were sectioned at 6 μm. Sections were incubated at 60°C for 1 h, then deparaffinized in xylene and rehydrated through graded ethanol to water. Antigen retrieval was omitted because preliminary optimization resulted in introducing nonspecific background without an apparent improvement in specific signal. IHC was performed using the ImmPRESS Excel Amplified Polymer Staining Kit (MP-7602-15, Vector Laboratories), according to the user guide with modifications. Endogenous peroxidase activity was quenched by incubating sections with BLOXALL Blocking Solution for 10 min at room temperature, followed by washing with PBS. Sections were then incubated with 2.5% normal horse serum for 20 min at room temperature to reduce nonspecific antibody binding. Without washing, sections were incubated overnight at 4°C with a mouse-derived anti-SARS-CoV-2 spike polyclonal antibody (75870-168, VWR INTERNATIONAL) diluted 1:10,000 in 1% BSA. The following day, sections were washed with PBS, incubated with the Amplifier antibody for 15 min at room temperature, washed with PBS, and then incubated with the ImmPRESS Polymer Reagent for 30 min at room temperature. After a PBS wash, immunoreactivity was visualized using ImmPACT DAB EqV substrate diluted to 1:10 in PBS for 2 min. The reaction was terminated by washing with PBS. Sections were counterstained with Harris modified hematoxylin diluted 1:1 with water, rinsed under running distilled water for 1.5 min, dehydrated through graded ethanol, cleared in xylene, and then mounted.

### 4. Metagenomic next-generation sequencing (mNGS)

#### 4.1 Library preparation and sequencing

Simultaneous reverse transcription of purified RNA, containing added (“spiked”) ERCC RNA internal controls (cat # 4456740, Invitrogen, Waltham, MA), and ribosomal RNA (rRNA) depletion were carried out using the NEBNext® Ultra™ II RNA First Strand Synthesis Module (cat #s E7771S/ E7771L, New England Biolabs, Ipswich, MA) and QIAseq FastSelect-rRNA HMR Kit (cat # 334385, Qiagen, Germantown, MD), respectively, followed by second strand cDNA synthesis using Sequenase™ Version 2.0 DNA Polymerase (cat # 70775Z1000UN, Thermo Fisher Scientific, Waltham, MA) as previously described^19^. The positive control for each run consisted of a previously described mixture of six representative organisms, including cytomegalovirus, spiked into buffer (Miller, et al., 2019). The negative control consisted of buffer spiked with T1 bacteriophage (DNA) and MS2 bacteriophage (RNA), which served as internal controls to verify detection of DNA and RNA viruses, respectively. The negative control consisted of T1 and M2 phages spiked into buffer. Complementary DNA (cDNA) was purified using NEBNext® Sample Purification Beads (cat # E7767L, New England Biolabs, Ipswich, MA) and end-repair was performed using the NEBNext® Ultra™ II RNA End Prep Module (cat #s E7771S/ E7771L, New England Biolabs, Ipswich, MA). Adaptor ligation, barcoding, and amplification (25 cycles) were performed using the NEBNext® Multiplex Oligos for Illumina kit (cat #’s E6440S/E6442S/E6444S/E6446S/ E6448S, New England Biolabs, Ipswich, MA) with additional cleanups also using NEBNext® Sample Purification Beads. Libraries were quantified and normalized using the Qubit dsDNA HS Assay (cat # Q32854, Thermo Fisher Scientific, Waltham, MA) on the Qubit Flex spectrophotometer (Thermo Fisher Scientific, Waltham, MA). Viral probe capture was carried out on separate normalized library pools using Twist Biosciences Standard Hybridization Solution v2 (SKU 104446, Twist Biosciences, South San Francisco, CA) and Comprehensive Viral Research Panel (SKU 103547, Twist Biosciences, South San Francisco, CA). DNA concentration was performed using the Alternate Pre-Hybridization DNA Concentration Protocol, which utilizes Twist DNA purification beads instead of a vacuum centrifuge. Final pooled libraries were sequenced as single-end reads on either the Illumina (San Diego, CA) NextSeq 550 using the Mid-Output or High-Output Kit (150 cycles) or the Illumina NovaSeq X using the 10B flow cell (300 cycles) to target depths of 1 million and 10-20 million reads per sample for libraries with and without viral probe capture, respectively.

#### 4.2 Bioinformatics

The SURPI+ computational pipeline, run as a container (research configuration, v1.0.13) on a secure local server, was used for agnostic identification of viral sequences from mNGS data. Reads were preprocessed by adapter trimming and removal of low-complexity and low-quality sequences, followed by computational subtraction of human reads using the Scalable Nucleotide Alignment Program^58^ (SNAP) nucleotide aligner with a maximum edit distance of 6. The SNAP aligner was then used to identify viral reads by aligning the remaining reads against microbial reference sequences in the National Center for Biotechnology Information (NCBI) nucleotide (NT) database (March 2019, with inclusion of the SARS-CoV-2 WuHan-Hu-1 genome) with a maximum edit distance of 16, followed by filtering and taxonomic classification of candidate viral hits^34^. Coverage maps were generated by automated mapping of SURPI+ -classified viral reads to the most likely reference genome. To identify sequence-divergent viruses, the pipelines used SPAdes (v3.15.4)^59^ and DIAMOND (v2.0.15)^60^, for de novo assembly of reads into contiguous sequences (contigs) and translated nucleotide sequence alignment for identification of sequence-divergent viruses, respectively. Reads from potential sequence-divergent viruses were identified uding DIAMOND alignment against reference viral sequences in the NCBI non-redundant (NR) protein database at an e-value cutoff of 10^−1^, with candidate viral hits filtered using a second SIAMOND search against the full NR protein database at an e-value cutoff of 10^−1^. No sequence-divergent viruses were detected in any of the study samples.

Quality control metrics for the mNGS libraries were based on those previously established for cerebrospinal fluid^19^, and include a minimum of 5 million preprocessed reads per sample, >75% of data with quality score >30 (Q > 30), and successful detection of the 6 organisms, including cytomegalovirus, in the positive control sample run in parallel. A positive viral detection was defined as the presence of ≥3 non-overlapping viral reads or contigs aligning to the target viral genome.

We used a customized nCounter Host Immune Response RNA expression panel (nanoString) to determine differential expression of 807 human host-viral response transcripts and viral transcripts across multiple genomic regions of SARS-CoV-2, SARS-CoV-1, and Epstein-Barr Virus (EBV). Pre-COVID-19 rectosigmoid tissue samples were used to identify false-positive signal and determine thresholds of positivity of SARS-CoV-2 transcript quantitation in the post-COVID samples.

### 5. nanoString nCounter targeted transcriptomics

nanoString ROSALIND software was used to perform differential gene expression analyses between LC and recovered participants. This approach selects the optimal subset of housekeeping probes for normalization using the geNorm algorithm, as implemented in the Bioconductor package. Differential expression between two groups of samples was calculated using the Generalized Linear Model (GLM) for count data assuming a negative binomial distribution and utilized the raw data and estimates of noise and dispersion across all samples in the experiment to calculate fold-change and p-value for each gene. Prior to DEG analyses, ROSALIND was used to remove genes/probes from differential expression analysis if they were expressed near or at background levels and then to normalize raw data. First, geometric mean of the selected housekeeping genes was calculated for each sample followed by geometric mean calculations across all sample-specific geometric means to generate a global geometric mean. Next, the global geometric mean was divided by the sample-specific geometric mean for each sample to generate a normalization factor for each sample followed by raw counts multiplied by of each gene by its sample-specific normalization factor.

### 6. Real-time PCR targeting the SARS-CoV-2 S1, and N regions

Quantitative PCR assays were performed using Integrated DNA Technology’s (IDT) SARS-CoV-2 RUO qPCR Primer & Probe Kits for S1, N, and RDRP (housekeeping transcript) detection (catalog no. 10006713, 10006804, 10006805, and 10006806). Positive controls consisted of fragments of human RPP30 and SARS-CoV-2 isolate Wuhan-Hu-1 (GenBank: NC_045512.2) provided in each IDT kit.

Quantitative PCR was performed with TaqPath 1-Step RT-qPCR Master Mix with the following conditions: 95 °C 2 min, 95 °C 3 sec, 55 °C 30 sec using the StepOnePlus Real-Time PCR System. SARS-CoV-2 RNA was considered detectable for cycle threshold (ct) values <40. Positive and non-template controls were run for all samples tested. Each sample was run in duplicate wells. Spike qPCR was also performed on RNA extracted from cryopreserved rectal tissue pieces using Thermo Fisher’s single-tube TaqMan Microbe Detection SARS-CoV-2 S gene Assay (catalogue no. A50137, Assay ID Vi07918636_s1) to detect the S1 domain of the spike protein. Thermo Fisher’s Comprehensive Microbiota Control (catalogue no. A50832), a multi-target plasmid pool containing the sequences for the S gene-specific assay, was used as a positive control. Quantitative PCR was run using TaqPath 1-Step RT-qPCR Master Mix with the following conditions: 50°C 2 min, 95°C 20 sec, 95°C 1 sec, 60°C 20 sec using the StepOnePlus Real-Time PCR System. SARS-CoV-2 RNA was considered detectable for cycle threshold (ct) values <40. Positive controls were run for all samples tested.

### 7. Meso Scale Discovery (MSD) antigen detection

Electrochemiluminescence immunoassays (S-PLEX SARS-CoV-2 Spike Kit & S-PLEX SARS-CoV-2 N Kit) were utilized to measure SARS-CoV-2 Spike and SARS-CoV-2 nucleocapsid in EDTA plasma from LIINC study participants. These S-PLEX assays have similar Recommended Protocols: (I) incubating (1 hour) the biotinylated antibody in the washed (three times, 150µL/well, gently tap all sides) S-PLEX 96-Well SECTOR or QuickPlex Plates that are streptavidin-coated, (II) incubating (1.5 hours) an equal volume (25µL) of sample/calibrator and Blocking Solution after washing the plate, (III) adding the TURBO-BOOST detection antibody solution after washing the plate then incubating (1 hour), (IV) incubating (30 mins) the washed plate with the MSD enhance solution, and (V) incubating (1 hour) the washed (three times, 150µL/well, gently tap all sides) plate with the TURBO-TAG detection solution. (VI) To detect the TURBO-TAG solution, 150µL/well of MSD GOLD Read Buffer B was added to a washed (three times) plate and read on an MSD reader (MESO QuickPlex SQ instrument). All incubations were performed on a plate shaker (700 rpm) at room temperature, with the exception of the TURBO-TAG incubation, which was completed on a Jitterbug at 27°C. For the N Kit, the Calibrator dilutions in (II) were prepared using Diluent 61 instead of Blocking Solution. All washes (performed manually) consisted of 2 plate washes with 150 µL/well per wash, followed by gentle tapping.

Spike positivity was defined using an empirical background-based threshold on the MSD assay. The maximum calculated concentration among samples classified as below the detection range was used to estimate background signal. Samples were considered positive if their calculated concentration exceeded five times this background value (≥5 × the maximum below-detection-range concentration).

### 8. Bulk-RNA sequencing

#### 8.1 Library and data preparation

RNA libraries, next-generation Illumina sequencing, quality control analysis, trimming and alignment were performed by Genewiz (Azenta). Briefly, following oligo dT enrichment, fragmentation and random priming, cDNA syntheses were completed. End repair, 5′ phosphorylation and dA-tailing were performed, followed by adaptor ligation, PCR enrichment and sequencing on an Illumina HiSeq platform using PE150 (paired-end sequencing, 150 bp for reads 1 and 2). Raw reads were trimmed using Trimmomatic (version 0.36) to remove adapter sequences and poor-quality reads. Trimmed reads were mapped to *Homo sapiens* GRCh37 using star aligner (version 2.5.2b)^61^.

#### 8.2 Data processing

Data was processed using nf-core/rnaseq v3.19.0^62^ of the nf-core collection of workflows^63^, utilizing reproducible software environments from the Bioconda^64^ and Biocontainers^65^. The pipeline was executed with Nextflow v25.04.6^63^. The sequence alignments were performed to a custom reference made up of human genome build hg38, HIV HIV1B and SARS-CoV-2 2018_SARS-CoV-2-wuhan-hu-1 sequences. The data were processed in two batches with a subset of samples in the second batch derived from the first batch. These repeated samples were reprocessed from the library preparation stage. Trim-galore^66^ was used to remove contaminated adapters and low-quality regions and the sequence quality was calculated and controlled with FastQC^67^. The sequencing reads were aligned to the custom reference using STAR^61^, the transcript levels quantified using Salmon^68^ and summarized to gene-level counts using tximport^69^.

Raw gene counts were assembled into a DGEList object using edgeR^70^ with associated gene and sample annotation. Low-expression genes were removed, retaining only genes with counts per million (CPM) > 0.5 in at least 2 samples. Log2-CPM values were computed for the full filtered dataset, and principal component analysis (PCA) was performed on the 2,000 most variable genes (by row variance) to assess overall sample structure, batch effects, and outliers. Samples with extreme PC1 scores (< −10) were flagged and removed as outliers. PCA was regenerated on the outlier-excluded dataset and visualized by sequencing batch, matched-sample identity, DNA/RNA extraction batch, self-reported race/ethnicity, and sex to screen for technical or demographic confounding. Samples separated by their processing batch on the PCA plot.

Batch-associated technical variation was estimated and corrected using RUVSeq^71^. Raw filtered counts were loaded into a SeqExpressionSet, upper-quartile between-lane normalization was applied. A set of empirical negative-control genes (the 2,000 genes with the lowest mean coefficient of variation, averaged across the two batches) was used to estimate one factor of unwanted variation (RUVg, k = 1). PCA plots before and after RUVSeq correction were generated and colored by batch, matched-sample identity, sex, race/ethnicity, and extraction batch to confirm that batch-associated structure was reduced with this choice of one factor of unwanted variation.

#### 8.3 Differential gene expression and gene set enrichment analysis

Differential gene expression was tested using edgeR’s quasi-likelihood negative binomial generalized linear model (GLM) framework, implemented per pairwise condition comparison. The design matrix modeled condition, the RUVSeq-derived factor of unwanted variation as a covariate, and sex. Differentially expressed genes (DEGs) were defined using an FDR threshold of 0.05 and an absolute log2 fold-change threshold of 0.25 (or 1, for selected downstream visualizations). Volcano plots showing the results of these association analyses were generated using the EnhancedVolcano bioconductor package^72^.

For comparisons yielding more than 10 DEGs (FDR < 0.05, |log2FC| > 0.25), functional enrichment was performed using clusterProfiler^73^ using Over-representation analysis (ORA) and Gene set enrichment analysis (GSEA). Gene symbols were mapped to Entrez IDs. DEGs (filtered at the same FDR and fold-change thresholds) were tested for enrichment using against custom term-to-gene mapping databases (GO^74^, Hallmark gene sets^75,76^) and KEGG pathways^77–79^ using the set of all expressed genes (non-zero counts across samples) as the statistical background/universe. All tested genes were ranked by a signed statistic (sign of log2 fold change × −log10[nominal p-value]) and analyzed with clusterProfiler::GSEA against the same term databases (minimum gene set size 5, maximum 500). Enrichment results were exported as tables (CSV) and visualized as bar plots, dot plots, and (for GSEA) ridge plots per database, with pathway descriptions truncated for display.

Batch-corrected normalized counts (RUVSeq output, log2-transformed) were mean-centered per gene and Winsorized at ±1 for visualization. The union of DEGs (FDR < 0.05) across all pairwise comparisons was used to define a gene set for heatmap display. Genes and samples were independently clustered by k-means (row k = 8, column k = 5 for the full DEG heatmap; row/column k = 3 for a top-N DEG heatmap restricted to the top 50 genes per comparison, ranked by p-value and filtered at FDR < 0.05 and |log2FC| > 1), and results were visualized as static heatmaps (using pheatmap^80^) ordered by cluster membership, annotated by symptom-category membership, using a blue–white–red diverging color scale. Gene clusters from the k-means row clustering were additionally each submitted to ORA (as described above) to identify cluster-specific enriched pathways.

Using the RUV-normalized, log2-transformed expression matrix restricted to a comparison of interest (LC≥1 vs. recovered), a signed gene co-expression network was constructed using the WGCNA package^81^. Genes and samples were quality-filtered using goodSamplesGenes. A soft-thresholding power was selected via pickSoftThreshold (powers 1–20, signed network) to approximate scale-free topology, and hierarchical clustering of samples (average linkage, Euclidean distance) was used to inspect for additional outliers. Network construction and module detection were performed using blockwiseModules (signed TOM, minimum module size = 30, module merge height = 0.35, maximum block size = 8,000 genes).

Module eigengenes were computed (moduleEigengenes, ordered via orderMEs) and merged with sample metadata (condition, sex, batch). Differential module expression between conditions was tested using limma (lmFit/contrasts.fit/eBayes on module eigengenes), with contrasts and BH-adjusted p-values reported per module. Gene-module membership (kME, via signedKME) was used to rank genes within each module, and gene set enrichment analysis was performed against Hallmark, GO, WikiPathways, and PFOCR gene sets using fgsea^82^, with Ensembl-to-gene-symbol mapping via org.Hs.eg.db.

### 9. OLINK plasma proteomics

Plasma proteomics was performed using the Olink® Reveal platform, which uses next-generation sequencing (NGS) following their proximity extension assay (PEA). The workflow was completed according to the Olink™ Reveal Laboratory Instructions. The plasma samples and controls (4µL each) were diluted with Sample Buffer and incubated overnight (19 hours) in the Reveal Incubation Plate for antibody binding.

The PCR Mix is then added to the Reveal Incubation Plate and placed into the thermocycler with the respective PCR Protocol initiated. These steps hybridize the oligonucleotides attached to the antibodies, allowing a unique sequence to be generated for each target protein. The library was purified using AMPure XP magnetic beads (Beckman Coulter). After illumina sequencing, ngs2counts (v6.2.0, Olink Proteomics) was used to generate protein levels expressed as NPX values, and NPX™ Map software (Olink Proteomics) was used to for plate normalization.

Normalized data were analyzed in R version 4.5.2. Differential protein abundance was assessed separately for each clinical phenotype (LC1, LC5, fatigue, taste/smell, neurologic symptoms, cardiopulmonary symptoms, gastrointestinal symptoms, and post-exertional malaise) by comparing phenotype-positive participants with recovered (R) controls. Proteins with insufficient data (<5 non-missing observations) were excluded before analysis. Differential abundance was evaluated using linear models with empirical Bayes moderation implemented in limma^83^, and p values were adjusted for multiple testing using the Benjamini-Hochberg false discovery rate (FDR) method. To identify enriched biological pathways, proteins were ranked by signed statistical significance (sign[log_2_ fold change] × −log_10_[P value]) and subjected to preranked Gene Set Enrichment Analysis (GSEA) using clusterProfiler^84^ and the MSigDB^85^ Hallmark gene sets.

### 10. Single cell RNA- and T cell receptor- (TCR) sequencing

#### 10.1 Library Preparation and data processing

Droplet-based scRNA-seq libraries were prepared using the 10x GEM-X Single Cell 5’ GEX and TCR for 16 frozen samples according to the manufacturer’s protocol. Libraries were sequenced at the Gladstone Institute Genomic Core using an Illumina NextSeq platform.

The fastq files were aligned to the 10x Genomics a customized version of the pre-built GRCh38 reference genome including HIV and SARS-CoV-2 genes using the 10x Genomics Cell Ranger v.8.0.0 count pipeline including introns^86^. The 10x Genomics Cell Ranger v.8.0.0 vdj pipeline (chain = TR) was used to reconstructs full-length V(D)J immune receptor sequences (contigs) for each TRA and TRB chain from each individual immune cell for downstream integration and visualization with gene counts. The raw count matrices generated by the Cell Ranger count pipeline were used as input into CellBender^87^ to adjust counts for ambient RNA contamination; expected cells were chosen for each sample after inspecting the barcode rank plots. The R package DropletUtils v1.20.0^88,89^ barcodeRanks function was used to identify the number of expected cells and the emptyDrops function was used to identify the empty droplet plateau. Following ambient RNA removal, each filtered count matrix was pre-processed as a Seurat v5.2.1^90^ object. The top 1% of cells per sample with a high number of unique genes and cells with ≤200 unique genes. After observing a high proportion of mitochondrial genes, the cells with ≥25% mitochondrial genes were filtered out. DoubletFinder^91^ was run with expected doublet formation rate to account for droplets containing more than one cell. Post quality control, individual Seurat objects were merged into a single Seurat object.

Normalization and variance stabilization were performed using sctransform v2^92^ selecting 15 principal components. Biological and technical variables were visualized using Seurat’s DimPlot function to assess the presence of batch effects in UMAPs. No cellular barcodes showing HIV and/or SARS-CoV-2 counts > 0 were detected.

#### 10.2 Clustering and cell type annotation

After excluding the highly expressed mitochondrial and TRC genes from the Variables Features list, the optimal clustering resolution parameters were determined using Random Forests^93^ and a silhouette score-based assessment of clustering validity and subject-wise cross-validation across multiple resolution parameters (0.02–1.2). This procedure is described in greater detail by George et al. 2022 and implemented in the clustOpt R package^94^. Graph-based clustering was performed using the Seurat v5.2.1 functions FindNeighbors and FindClusters using the Louvain algorithm setting the optimal resolution to 0.08 and using 15 principal components resulted in 13 distinct clusters. Marker genes for each cluster were identified using Seurat’s FindAllMarkers function with the Wilcoxon rank-sum test (minimum percentage of cells expressing a marker: 25%) Cell types were assigned to each cluster by providing the FindAllMarkers results for each cluster to cellKB online tool [ref 10], with gut wall and gut lymphoid tissue-specific genes sets. After annotation, sample 7425 specific-cluster 5 was excluded from the downstream analyses.

#### 10.3 Cell composition association with phenotype

Associations between cell type composition and phenotype (“Pre-COVID-19”,““Long-COVID”, “Recovered”) were assessed using the LODopt package^95^. LODopt estimates changes in absolute cell type composition while accounting for the compositional nature of single-cell data through log-odds ratio optimization. Cell counts were aggregated by cluster and sample. The method uses a generalized linear mixed-effects model (GLMM) framework with phenotype “Long-COVID” as the predictor variable and “Recovered” as the reference level. Normalization factors were optimized to minimize bias in log-odds ratio estimates.

Statistical significance was assessed at p < 0.05.

#### 10.4 TCR and scRNAseq data integration

The sample-specific filtered contig annotations were integrated in Seurat object using combine TCR function implemented in scRepertoire^96^. The 2025 list of SARS-CoV-2 specific epitope were downloaded from VDJdb [vdjdb.cdr3.net]. Cellular barcodes expressing low to high confidence SARS-CoV-2 CDR3 sequences (score >= 1) and high confidence (score > 2) were tagged in the Seurat object. Dimensional plots showing the distribution of entire repertoire of TCR, the low to high and high confidence SARS-CoV-2 TCR by phenotype were generated.

#### 10.5 Pseudobulk differential expression (DE)

Pseudobulk DE analysis was performed using the muscat R package^97^. Raw gene counts were aggregated across cells within each sample and cluster to generate a pseudo-bulked expression matrix. Only clusters with a median of at least 10 cells per sample were retained for downstream analysis. Differential expression was evaluated for phenotype as the sole predictor using the pbDS function from muscat with the edgeR method^70^. Design matrices included phenotype as the predictor variable. The comparisons of interest were (1) “Pre-COVID” vs ‘Post-COVID” (combining “Long-COVID” and “Recovered”), and (2) “Long COVID” vs “Recovered”. The min_cells parameter in pbDS was set to 3 to ensure that each pseudobulk sample contained at least three cells. The pvalue were adjusted for multiple test comparison using BH method. The Seurat object was subset to extract the list of nCounter genes. The analyses was performed as above and adjusted for multiple comparisons. Differentially expressed genes were visualized using EnhancedVolcano R package^72^.

#### 10.6 Gene set enrichment analysis (GSEA)

Differentially expressed genes (p-value < 0.0.5 and absolute log₂ fold change > 0.25) from the two comparisons were used for over-representation analysis (ORA) using the clusterProfiler package^73^. For GSEA, all genes were ranked by sign(log2FC) × –log₁₀(p-value) without filtering. Gene symbols were mapped to Entrez IDs using the bitr function from clusterProfiler. Enrichment analyses were performed using Gene Ontology Biological Processes^74,98^, KEGG^99^, WikiPathways^100^, and PFOCR^101^ databases with minimum and maximum gene set sizes of 5 and 500, respectively

### 11. 27-plex Flow Cytometry

The phenotype of GALT cells was characterized by flow cytometry using Cytek 25-plex immunophenotyping panel (Cytek Biosciences) with a modification by adding goat F(ab’)2 anti-human IgA-Biotin (Southern Biotech) and BUV395 Streptavidin (BD Biosciences). During the panel development, each antibody utilized in the panel was titrated using PBMC to optimize concentration. Fluorescent minus one (FMO) staining for IgA was used to determine baseline expression level.

GALT cells were stained with Zombie UV dye in PBS for 10 minutes at 25°C for live-cell gating followed by incubation with True-Stain Monocyte Blocker^TM^ (Biolegend) for 10 min before adding surface staining markers in PBS+1%FBS with Brilliant Stain Buffer Plus (BD Biosciences) in a total volume of 75 ml for 20 minutes at 25°C according to the manufacturer’s protocol. Following surface staining, samples were resuspended in PBS+1%PFA and stored at 4°C before acquiring on a Cytek 5L Aurora (Cytek Biosciences, Fremont, CA, USA) using SpectroFlo (v3.0.3). Compbead anti-mouse, -rat and -hamster particles (BD Biosciences) were stained as reference controls for unmixing except that heat-killed PBMCs were used as a Zombie UV staining unmixing reference. For each patient sample, one unstained control tube was prepared for autofluorescence reduction.

Prior to data acquisition, instrument setup and quality control (QC) procedures were performed according to the manufacturer’s recommendations. Data were acquired using the Cytek 25-color Immunoprofiling Assay (Cytek DOC-00300 Rev. B). Approximately 0.5 × 10 events were collected per sample. Single-stained BD™ CompBeads and PBMC controls were acquired with the unmixing matrix, and autofluorescence extraction was performed using SpectroFlo (v3.0.3) with the recommended reference controls (**Table S9**). Multicolor samples were subsequently acquired using the live unmixing function. Spillover adjustments were applied, as needed, to correct unmixing errors according to Cytek’s recommendations.

Flow cytometry data were analyzed using FlowJo software (v10.10.1; BD Life Sciences). Gating strategies are shown in **Fig. S6-8**. Population frequencies were exported for downstream statistical analysis and visualization in GraphPad Prism.

### 12. Xenium Digital Spatial Profiling (DSP)

#### 12.1 Library preparation

10X Genomics Xenium In-Situ Gene Expression assay was performed according to the manufacturer’s protocol using a human pan-tissue commercial base panel with 100 additional human and viral transcripts including extended colorectal tissue- and immune-related transcripts including SARS-CoV-2 RNA, EBV and CMV as in **Table S8**. Briefly, 5 um FFPE sections were mounted on Xenium slides and deparaffinized and decrosslinked. Next, slides were incubated overnight with the Xenium Human Colon Gene Expression Panel (10X Genomics, 1000642) and a custom add-on gene panel. Following probe hybridization, slides were treated with a post-hybridization wash, followed by ligation, amplification, and a post-amplification wash. Next, slides were incubated overnight with the multi-tissue cell segmentation stain mix. Finally, autofluorescence quenching and nuclei staining were performed. Following assay completion, slides were transferred to PBS-T and immediately scanned on the Xenium Analyzer. H&E staining was performed after Xenium Analyzer run.

#### 12.2 Cell segmentation

Individual cell segmentation was determined using microscopic images of nuclear material, cytoplasmic RNA and cell wall staining from 5 µm tissue sections on the Xenium Explorer platform. Transcript counts from the Xenium run and spatial coordinates were then exported for further analysis in Python

#### 12.3 Clustering

To generate robust and reproducible cell clusters, we employed negative binomial count splitting to partition the raw count matrix into statistically independent training and testing datasets, allowing cluster stability to be evaluated without introducing information leakage between model training and validation^102^.

Gene expression counts were modeled as:

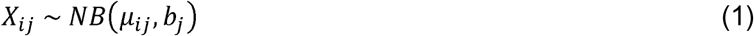

where *b_j_* represents the gene-specific dispersion parameter. Independent count matrices were generated through Dirichlet–Multinomial thinning, such that

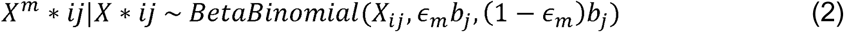

Because the true dispersion parameters were unknown, they were estimated for each gene using the method of moments,

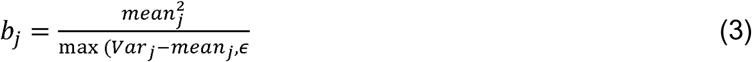

The raw counts were then partitioned into 60% training data and 40% testing data. Given the large dataset size (277,202 cells), this allocation provided sufficient training data for model fitting while preserving an independent test set for validation.

The training data were used to fit an scVI model for dimensionality reduction and latent representation learning^103^. The model consisted of two hidden layers with a 30-dimensional latent space and incorporated plate information to correct for batch effects. A nearest-neighbor graph was then constructed using a neighborhood size of 40, followed by UMAP for two-dimensional visualization^104–106^. The testing data were projected into the pretrained scVI latent space without further model training.

Leiden clustering was performed independently on both the training and testing latent representations across multiple resolution parameters^107–109^. Cluster agreement between the corresponding training and testing cells was evaluated using the Adjusted Rand Index (ARI). The optimal clustering was obtained at a Leiden resolution of 0.32 (ARI = 0.344). This parameter was subsequently applied to the complete dataset, resulting in the final set of nine clusters.

#### 12.4 Cell type annotation

Cell type annotation was performed using the clusters identified in the previous analysis. Cluster identities were assigned by integrating multiple sources of evidence, including spatial localization, scVI differential expression analysis, Scanpy differential expression rankings, and cluster-specific marker genes.

#### 12.5 Pathway enrichment activity scores

Pathway enrichment analysis was performed to identify biological pathways differentially represented between LC and control groups within each annotated cell type.

Gene enrichment analysis was performed using the decoupler Python package on cell-area-normalized, log-transformed data^110^. Pathway activities were inferred using the Univariate Linear Model (ULM) method with PROGENy, MSigDB, KEGG, Hallmark, and CytoSig gene sets [10–12]^76,111,112^.

Pathways with fewer than five overlapping gene targets were excluded. Because the dataset contained only 477 measured genes, pathway genes were not weighted according to lfcmean. Pathway activities were ranked using a t-test with overestimated variance, producing adjusted p-values and mean-change statistics.

