## Supplemental Tables and Figures for "Multiomic and Spatial Profiling of Colorectal Tissue Reveals Viral Persistence and Immune Dysregulation in Long COVID"

### Table S1: List of all 807 genes included on nCounter panel

| | | | | | | | | |
|---|---|---|---|---|---|---|---|---|
| ACE | CCL17 | CTSL | GBP1 | IKBKE | IRF4 | MS4A4A | PSMB9 | TAB2 |
| ACE2_Hs | CCL18 | CTSS | GBP2 | IKBKG | IRF7 | MS4A7 | PSTPIP1 | TANK |
| ACKR2 | CCL19 | CTSW | GBP4 | IL10 | IRF9 | MSRA | PTGER2 | TAP1 |
| ACKR3 | CCL2 | CTSZ | GBP5 | IL10RA | ISG15 | MT2A | PTGER4 | TAP2 |
| ACKR4 | CCL20 | CUL1 | GCA | IL10RB | ITGAE | MTOR | PTGS2 | TBK1 |
| ACOX1 | CCL21 | CX3CL1 | GK | IL11 | ITGAL | MVP | PTK2B | TBX21 |
| ACSL1 | CCL22 | CX3CR1 | GLA | IL11RA | ITGAM | MX1 | PTPN4 | TBXAS1 |
| ACSL3 | CCL23 | CXCL1 | GLB1 | IL12A | ITGAX | MYC | PTPN6 | TCF7 |
| ACSL4 | CCL24 | CXCL10 | GNLY | IL12B | ITGB2 | MYD88 | PTPRC | TCIRG1 |
| ACVR1 | CCL25 | CXCL11 | GNS | IL12RB1 | ITGB7 | NAE1 | PXN | TCL1A |
| ADAR | CCL26 | CXCL12 | GPX7 | IL12RB2 | ITK | NAMPT | PYCARD | TCN2 |
| ADGRE5 | CCL27 | CXCL13 | GSK3B | IL13 | ITLN1 | NCF1 | RAB31 | TGFB1 |
| ADGRG3 | CCL28 | CXCL14 | GSTM4 | IL13RA1 | ITPR3 | NCF2 | RAB5C | TGFB2 |
| ADORA2A | CCL3/L1/L3 | CXCL16 | GUCY1A1 | IL13RA2 | JAK1 | NCF4 | RAB7A | TGFB3 |
| AGT | CCL4/L1/L2 | CXCL17 | GUCY1B1 | IL15 | JAK2 | NCR1 | RAC2 | TGFBR2 |
| AHR | CCL5 | CXCL2 | GZMA | IL15RA | JAK3 | NCR3 | RACK1 | THBS1 |
| AIF1 | CCL7 | CXCL3 | GZMB | IL16 | JAML | NDUFS8 | RAF1 | THOP1 |
| AIM2 | CCL8 | CXCL5 | GZMH | IL17A | JUN | NEO1 | RASGRP1 | TIFA |
| AKT1 | CCNC | CXCL6 | HAMP | IL17B | JUNB | NEU1 | RASGRP4 | TIGIT |
| AKT2 | CCR1 | CXCL8 | HAVCR2 | IL17C | KDM6B | NFAT5 | RB1CC1 | TIMP2 |
| AKT3 | CCR10 | CXCL9 | HCK | IL17D | KIR2DL1 | NFATC1 | RBCK1 | TLN1 |
| ALOX12 | CCR2 | CXCR1 | HCoV-229E_N | IL17F | KIR2DL3 | NFATC2 | RBPJ | TLR1 |
| ALOX15 | CCR3 | CXCR2 | HCoV-229E_S | IL17RA | KIR3DL1/2 | NFATC3 | REL | TLR2 |
| ALOX5 | CCR4 | CXCR3 | HCoV-HKU1_N | IL17RB | KLRB1 | NFATC4 | RELA | TLR3 |
| ALOX5AP | CCR5 | CXCR4 | HCoV-HKU1_S | IL17RC | KLRC1 | NFE2L2 | RELB | TLR4 |
| ALPK1 | CCR6 | CXCR5 | HCoV-NL63_N | IL17RD | KLRD1 | NFKB1 | RGMA | TLR5 |
| ALPL | CCR7 | CXCR6 | HCoV-NL63_S | IL17RE | KLRK1 | NFKB2 | RHOG | TLR6 |
| ANPEP | CCR8 | CYP2E1 | HCoV-OC43_N | IL18 | KPNB1 | NGLY1 | RIPK1 | TLR7 |
| AP1G1 | CCR9 | CYSTM1 | HCoV-OC43_S | IL18BP | KRAS | NKG7 | RIPK2 | TLR8 |
| AP1M1 | CCRL2 | DDAH2 | HCST | IL18R1 | LAG3 | NLRC4 | RIPK3 | TLR9 |
| AP1S2 | CD14 | DDIT3 | HDC | IL18RAP | LAMP1 | NLRC5 | RNASEL | TMEM140 |
| APBB1IP | CD163 | DDOST | HERC5 | IL19 | LAMP2 | NLRP1 | RNF114 | TMPRSS2 |
| APEX1 | CD19 | DDX5 | HK3 | IL1A | LAMP3 | NLRP3 | RNF135 | TNF |
| APOBEC3G | CD1E | DDX58 | HLA-A | IL1B | LANCL1 | NOD2 | RNF31 | TNFRSF10B |
| APOL6 | CD2 | DEFA4 | HLA-B | IL1F10 | LAT | NOS2 | RPS6KA1 | TNFRSF17 |
| APP | CD209 | DEFB103A/B | HLA-C | IL1R1 | LAT2 | NOTCH1 | RPS6KA3 | TNFRSF18 |
| ARRB2 | CD22 | DERL1 | HLA-DMA | IL1R2 | LCK | NOX1 | RPS6KB1 | TNFRSF1A |
| ATF2 | CD244 | DHX58 | HLA-DMB | IL1RAP | LCN2 | NPC2 | RSAD2 | TNFRSF25 |
| ATF4 | CD247 | DIABLO | HLA-DOB | IL1RAPL1 | LCP1 | NRAS | RUNX3 | TNFRSF4 |
| ATF6 | CD27 | DNAJA2 | HLA-DPA1 | IL1RAPL2 | LCP2 | NT5E | S100A12 | TNFRSF8 |
| ATG10 | CD274 | DNAJC10 | HLA-DPB1 | IL1RL1 | LDHB | NTNG2 | SAMHD1 | TNFRSF9 |
| ATG12 | CD276 | DTX3L | HLA-DQA | IL1RL2 | LEF1 | OAS1 | SARS-CoV_N | TNFSF10 |
| ATG13 | CD28 | DYSF | HLA-DQB1 | IL1RN | LGALS3 | OAS2 | SARS-CoV_S | TNFSF13B |
| ATG3 | CD36 | EBI3 | HLA-DRA | IL2 | LIF | OAS3 | SARS-CoV-2_E | TNFSF18 |
| ATG4A | CD38 | EBV_BLLF1 | HLA-DRB | IL20 | LILRA3 | OASL | SARS-CoV-2_M | TNFSF4 |
| ATG7 | CD3D | EBV_EBER1 | HLA-E | IL20RA | LILRA5 | OS9 | SARS-CoV-2_N | TNFSF9 |
| ATM | CD3E | EBV_EBER2 | HLX | IL20RB | LILRA6 | OSM | SARS-CoV-2_orf1ab | TOLLIP |
| ATP6AP2 | CD3G | EBV_EBNA1 | HMGB1 | IL21 | LILRB2 | P2RX7 | SARS-CoV-2_orf1ab_REV | TPP1 |
| ATP6V0D1 | CD4 | EBV_EBNA2_1 | HMOX1 | IL21R | LIMK2 | PAK1 | SARS-CoV-2_ORF3a | TPSAB1/B2 |
| ATP6V1B2 | CD40 | EBV_EBNA2_2 | HPGD | IL22 | LITAF | PANX1 | SARS-CoV-2_ORF7a | TRAF2 |
| BATF | CD40LG | EBV_EBNA3A | HSD11B1 | IL22RA1 | LRG1 | PARP1 | SARS-CoV-2_ORF8 | TRAF3 |
| BCL2 | CD44 | EBV_EBNA3BC | HSP90AA1 | IL22RA2 | LRRK2 | PARP9 | SARS-CoV-2_S | TRAF6 |
| BCL2L1 | CD45R0 | EBV_LMP1 | HSP90AB1 | IL23A | LTA4H | PDCD1 | SCARB2 | TRAM1 |
| BCL3 | CD45RA | EBV_LMP2A | HSP90B1 | IL23R | LTB | PDCD1LG2 | SELE | TRAT1 |
| BCL6 | CD45RB | EBV_LMP2A/B | ICAM3 | IL24 | LTBR | PDHB | SELENOS | TRIM21 |
| BCR | CD59 | EGLN1 | ICOS | IL25 | LTC4S | PECAM1 | SELL | TRIM22 |
| BDKRB1 | CD6 | EIF2AK2 | ICOSLG | IL26 | LTF | PELI1 | SEM1 | TRIM25 |
| BDKRB2 | CD68 | EIF2AK3 | IDO1 | IL27 | LYN | PELI2 | SERPINA1 | TRIM33 |
| BECN1 | CD69 | EIF3F | IFI16 | IL27RA | MAF | PFKFB3 | SH2D1A | TRIM5 |
| BLK | CD70 | ELANE | IFI27 | IL2RA | MAFB | PIK3C3 | SIGIRR | TRIM56 |
| BNIP3 | CD79A | ENTPD1 | IFI35 | IL2RB | MAP1LC3A | PIK3CA | SIGLEC5 | TRIM6 |
| BPI | CD79B | EOMES | IFI44 | IL2RG | MAP2K2 | PIK3CB | SIRPA | TXK |
| BST2 | CD80 | EPHX2 | IFI6 | IL3 | MAP2K3 | PIK3CD | SLC11A1 | TXN |
| C1QBP | CD81 | ERN1 | IFIH1 | IL31 | MAP2K4 | PIK3CG | SLC2A3 | TXNIP |
| C2 | CD84 | ETS1 | IFIT1 | IL31RA | MAP2K7 | PIK3R3 | SMAD3 | TYK2 |
| C3 | CD86 | EVL | IFIT2 | IL32 | MAP3K1 | PIK3R4 | SMAD4 | TYROBP |
| C3AR1 | CD8A | F5 | IFIT3 | IL33 | MAP3K3 | PIK3R5 | SMAD5 | UBA52 |

| | | | | | | | | |
|---|---|---|---|---|---|---|---|---|
| C5 | CD8B | FAM30A | IFITM1 | IL34 | MAP3K5 | PIK3R6 | SOCS1 | UBE2L6 |
| C5AR1 | CDH1 | FAS | IFITM2 | IL36A | MAP3K7 | PLAT | SOCS3 | UBE2N |
| CALM1 | CDK4 | FASLG | IFITM3 | IL36B | MAP3K8 | PLAU | SOD1 | ULK1 |
| CAP1 | CEACAM3 | FBXO6 | IFNA1/13 | IL36G | MAPK1 | PLAUR | SOD2 | ULK2 |
| CARD11 | CEBPB | FCAR | IFNA14/16 | IL36RN | MAPK13 | PLCG1 | SORT1 | VAMP3 |
| CARD16 | CFLAR | FCGR1A/B | IFNA2 | IL37 | MAPK14 | PLCG2 | SP1 | VCAM1 |
| CARD17 | CGAS | FCGR2A | IFNA4/7/10/17/21 | IL3RA | MAPK8 | PLEK | SP100 | VEGFA |
| CARD8 | CHUK | FCGR3A/B | IFNA5 | IL4 | MAPK9 | PLEKHA1 | SPI1 | VRK3 |
| CASP1 | CPA3 | FCGRT | IFNA6 | IL4R | MAPKAPK2 | PLG | SPIB | VSIR |
| CASP10 | CR1 | FCRL2 | IFNA8 | IL5 | MARCKS | PLIN4 | SSR1 | VWF |
| CASP3 | CREBBP | FCRL4 | IFNAR1 | IL5RA | MARCO | PNOC | STAT1 | WAS |
| CASP4 | CRK | FGR | IFNAR2 | IL6 | MAVS | PPIA | STAT2 | WIPI1 |
| CASP5 | CRP | FOS | IFNB1 | IL6R | MCL1 | PRCP | STAT3 | XAF1 |
| CASP8 | CSF1 | FOXO1 | IFNG | IL6ST | MDFIC | PRDM1 | STAT4 | XBP1 |
| CBFB | CSF1R | FOXP3 | IFNGR2 | IL7 | MEFV | PRF1 | STAT5A | XCL1/2 |
| CBL | CSF2 | FPR1 | IFNK | IL7R | MGAM | PRKCA | STAT5B | XCR1 |
| CBLB | CSF2RA | FPR2 | IFNL1 | IL9 | MIF | PRKCD | STAT6 | YWHAQ |
| CCL1 | CSF2RB | FURIN | IFNL2/3 | IL9R | MKNK1 | PRKCQ | STING1 | ZAP70 |
| CCL11 | CSF3 | FYN | IFNL4 | IRAK1 | MLKL | PRKCSH | STRAP | ZBP1 |
| CCL13 | CSF3R | GAB2 | IFNLR1 | IRAK3 | MME | PSAP | STT3B | |
| CCL14 | CTLA4 | GADD45B | IFNW1 | IRAK4 | MRC1 | PSEN1 | SUGT1 | |
| CCL15 | CTSA | GATA3 | IGFBP7 | IRF1 | MS4A1 | PSMB10 | SYK | |
| CCL16 | CTSG | GBA | IKBKB | IRF3 | MS4A2 | PSMB8 | TAB1 | |

**Table 2: Participant demographics for those with and without detection of SARS-CoV-2**

| Characteristic | Participants without SCV2 persistence N = 44[1] | Participants with SCV2 persistence N = 13[1] | p-value |
|---|---|---|---|
| **Age** (years) | 45 (37, 58) | 43 (41, 54) | >0.9 |
| **Sex** | | | 0.5 |
| Female | 28 (64%) | 7 (54%) | |
| Male | 16 (36%) | 6 (46%) | |
| **Race/Ethnicity** | | | 0.2 |
| American Indian or Alaska Native | 0 (0%) | 1 (7.7%) | |
| Asian | 6 (14%) | 0 (0%) | |
| Black/African American | 2 (4.5%) | 0 (0%) | |
| Hispanic/Latino | 4 (9.1%) | 2 (15%) | |
| White | 32 (73%) | 10 (77%) | |
| **Medical comorbidities** | | | |
| Autoimmune disease | 6 (14%) | 1 (7.7%) | >0.9 |
| Cancer[2] | 1 (2.3%) | 0 (0%) | >0.9 |
| Diabetes | 1 (2.3%) | 1 (7.7%) | 0.4 |
| Hypertension | 7 (16%) | 1 (7.7%) | 0.7 |
| Lung disease[3] | 8 (18%) | 4 (31%) | 0.4 |
| **Body mass index** (kg/m2) | | | 0.2 |
| <25 | 25 (57%) | 4 (31%) | |
| 25-29.9 | 12 (27%) | 6 (46%) | |
| >= 30 | 7 (16%) | 3 (23%) | |
| **Hospitalized for COVID-19** | 2 (4.5%) | 2 (15%) | 0.2 |
| **Antiviral treatment for acute COVID-19** | 6 (14%) | 3 (23%) | 0.4 |
| **First SVC2 infection era** | | | 0.4 |
| March 2020 – April 2021 | 9 (20%) | 5 (38%) | |
| May 2021 – November 2021 | 3 (6.8%) | 0 (0%) | |
| >December 2021 | 32 (73%) | 8 (62%) | |
| **Time since first SCV2 infection** (days) | 725 (505, 903) | 565 (275, 764) | 0.09 |
| **Time since most recent SCV2 infection** (days) | 575 (444, 809) | 275 (188, 728) | 0.07 |
| **Vaccinated prior to first SCV2 infection** | 34 (77%) | 8 (62%) | 0.3 |
| **Vaccinated prior to colorectal biopsy**[4] | 43 (98%) | 12 (92%) | >0.9 |
| **Quality of life by visual analog scale**[4,5] | 65 (50, 80) | 65 (60, 70) | 0.9 |
| **Long COVID symptom count**[4] | 8 (5, 10) | 6 (4, 12) | 0.2 |
| **Symptoms reported**[6] | | | |
| Fatigue | 29 (66%) | 10 (77%) | 0.6 |
| Cardiopulmonary | 13 (30%) | 5 (38%) | >0.9 |
| Gastrointestinal | 17 (39%) | 4 (31%) | 0.3 |
| Smell/Taste disturbance | 5 (11%) | 2 (15%) | >0.9 |
| Neurocognitive | 31 (70%) | 10 (77%) | 0.2 |
| Post-Exertional Malaise | 23 (52%) | 6 (46%) | 0.3 |
| Sleep disturbance | 21 (48%) | 6 (46%) | 0.5 |

[1]n (%); Median (IQR). [2]Cancer requiring treatment within the 2 years before COVID-19. [3]Asthma, COPD, emphysema or bronchitis experienced in the 5 years prior to COVID-19. [4]At time of biopsy. [5]Data missing for one participant.[6]**Cardiopulmonary:** cough, shortness of breath, heart palpitations, chest pain. **Gastrointestinal:** diarrhea, nausea, loss of appetite, abdominal pain, vomiting, constipation. **Neurocognitive:** concentration problems, dizziness, balance problems, neuropathy, vision problems.

**Table S3: List of all 240 Significant DEGs from nCounter targeted transcriptomics analysis in colorectal samples from people with LC vs recovered controls***

| | | | | | | | |
|---|---|---|---|---|---|---|---|
| CCL8 | RNF135 | HLA-DMB | TRAM1 | MGAM | BCL6 | FPR1 | IL1RL1 |
| CCL13 | STAT3 | CD276 | PLAUR | SOD2 | CD81 | CBL | CCL20 |
| NEU1 | HLA-B | CASP10 | EPHX2 | TNFRSF4 | LCN2 | PIK3R5 | ITGAM |
| IL13RA1 | CCR3 | SAMHD1 | CD163 | PRCP | SORT1 | FCGR2A | NAE1 |
| MRC1 | CCL11 | ALOX5AP | STING1 | VRK3 | HERC5 | TNFSF13B | FPR2 |
| CSF1R | GNS | CXCL16 | RB1CC1 | MAF | OS9 | FCGRT | CD244 |
| JAML | RNASEL | BCR | HLA-DMA | ENTPD1 | CBFB | GSK3B | IRF7 |
| LANCL1 | MAP2K3 | ITGB2 | NOX1 | CD209 | MARCKS | TRAF2 | CTSS |
| FCGR3A/B | PIK3CG | BCL3 | GBA | IFIH1 | MAPK14 | PIK3R4 | PIK3CA |
| TRIM25 | S100A12 | CTSA | GBP4 | SCARB2 | ALPK1 | ALOX5 | CTSZ |
| WIPI1 | TLR7 | ICAM3 | MAPK1 | TLR8 | STAT1 | CD44 | VWF |
| RBPJ | PSMB9 | NCF4 | PTPN4 | VAMP3 | TNFRSF25 | IL33 | CD14 |
| TAB2 | PAK1 | MAP3K1 | TNFRSF10B | HLA-DRA | CCL14 | TCN2 | DIABLO |
| IL1RN | RPS6KB1 | LTBR | ULK1 | ATP6AP2 | IRAK1 | UBA52 | CBLB |
| PSAP | TPP1 | HLA-E | CSF2RA | CRK | YWHAQ | UBE2N | SEM1 |
| TLR3 | IRF3 | IFI27 | STAT6 | PYCARD | FAS | LCP2 | HLA-DPA1 |
| TXNIP | NFKB2 | AIF1 | IL7 | IL22RA1 | PLCG2 | CSF1 | C1QBP |
| ACOX1 | DDAH2 | ATF2 | ADAR | VSIR | LAG3 | ATP6V1B2 | CD45R0 |
| SMAD3 | SP100 | CD3G | DDX5 | SPIB | HSP90AB1 | CD68 | CXCL11 |
| CD36 | ACKR3 | CXCL14 | GPX7 | KIR2DL3 | MAPKAPK2 | IL1F10 | TRAT1 |
| SMAD4 | DERL1 | LAMP2 | SOD1 | AKT2 | TRIM6 | PRKCD | FCAR |
| GZMA | MAPK9 | ITGAE | CD84 | ANPEP | HLA-DPB1 | ACSL3 | RPS6KA3 |
| ATG13 | MS4A4A | MLKL | IL10RA | EGLN1 | HLA-A | PRDM1 | PANX1 |
| IFNLR1 | NLRC4 | SERPINA1 | STT3B | KLRD1 | IFITM1 | ACKR4 | IL2RG |
| CCL24 | TLR5 | HSP90AA1 | BST2 | APOL6 | SELENOS | LRG1 | APP |
| CD4 | ADGRE5 | ATF4 | ITGB7 | ITLN1 | LAMP1 | MDFIC | TAP1 |
| PSMB8 | EIF3F | TLR1 | HAVCR2 | C3AR1 | MAP2K4 | CASP3 | NCR1 |
| HSP90B1 | MS4A7 | HMOX1 | CASP8 | NKG7 | TNFSF4 | CYSTM1 | EBV_EBNA3BC |
| LTA4H | PSMB10 | TMPRSS2 | CD59 | ATP6V0D1 | SELE | HCST | FURIN |
| PXN | CCL7 | MAP3K5 | HCK | HLA-DQA | TLR6 | CCL22 | IL18 |

*Pathways ordered from lowest to highest FDR adjusted p-value.

**Table S4: Significantly enriched gene sets from unbiased RNA sequencing in colorectal samples from participants with LC compared to recovered controls.**

| Pathway Name | setSize[1] | NES[2] | FDR Adjusted P-Value |
|---|---|---|---|
| **Hallmark Pathways** | | | |
| MYC TARGETS V2 | 58 | 2.53 | 1.25E-09 |
| MYC TARGETS V1 | 200 | 2.21 | 1.25E-09 |
| MTORC1 SIGNALING | 200 | 2.18 | 1.25E-09 |
| OXIDATIVE PHOSPHORYLATION | 199 | -2.15 | 1.25E-09 |
| UNFOLDED PROTEIN RESPONSE | 112 | 2.20 | 9.87E-09 |
| G2M CHECKPOINT | 199 | 1.94 | 4.74E-08 |
| PI3K AKT MTOR SIGNALING | 100 | 1.96 | 1.15E-05 |
| TNFA SIGNALING via NFKB | 198 | 1.62 | 6.28E-04 |
| E2F TARGETS | 200 | 1.57 | 8.04E-04 |
| CHOLESTEROL HOMEOSTASIS | 73 | 1.80 | 8.35E-04 |
| GLYCOLYSIS | 188 | 1.60 | 9.23E-04 |
| APICAL JUNCTION | 186 | 1.54 | 2.43E-03 |
| ESTROGEN RESPONSE EARLY | 195 | 1.53 | 2.88E-03 |
| UV RESPONSE UP | 152 | 1.55 | 6.32E-03 |
| ADIPOGENESIS | 195 | -1.47 | 6.32E-03 |
| INFLAMMATORY RESPONSE | 193 | 1.45 | 6.98E-03 |
| BILE ACID METABOLISM | 98 | -1.55 | 1.12E-02 |
| HEME METABOLISM | 186 | -1.43 | 1.27E-02 |
| ESTROGEN RESPONSE LATE | 195 | 1.41 | 1.27E-02 |
| PANCREAS BETA CELLS | 34 | 1.65 | 1.70E-02 |
| IL6 JAK STAT3 SIGNALING | 85 | 1.51 | 1.75E-02 |
| ANDROGEN RESPONSE | 100 | 1.45 | 2.27E-02 |
| COMPLEMENT | 188 | 1.38 | 2.45E-02 |
| MITOTIC SPINDLE | 199 | 1.34 | 2.49E-02 |
| TGF BETA SIGNALING | 54 | 1.51 | 2.77E-02 |
| PROTEIN SECRETION | 96 | 1.43 | 2.77E-02 |
| **KEGG Pathways** | | | |
| Ribosome | 172 | -2.99 | 5.90E-09 |
| Coronavirus disease | 203 | -2.69 | 5.90E-09 |
| Spliceosome | 141 | 2.49 | 5.90E-09 |
| Oxidative phosphorylation | 126 | -2.49 | 5.90E-09 |
| Ribosome biogenesis in eukaryotes | 75 | 2.39 | 5.90E-09 |
| Protein processing in endoplasmic reticulum | 165 | 2.24 | 5.90E-09 |
| Nucleocytoplasmic transport | 102 | 2.26 | 1.08E-08 |
| Diabetic cardiomyopathy | 193 | -2.07 | 1.37E-08 |
| Cornified envelope formation | 116 | 2.15 | 7.58E-08 |
| Pathogenic Escherichia coli infection | 182 | 1.98 | 3.77E-07 |
| TNF signaling pathway | 117 | 2.07 | 1.91E-06 |
| Cadherin signaling | 289 | 1.79 | 1.91E-06 |
| Salmonella infection | 238 | 1.82 | 4.39E-06 |
| mRNA surveillance pathway | 85 | 2.08 | 9.35E-06 |
| Hepatitis C | 128 | 1.91 | 1.59E-05 |
| Bacterial invasion of epithelial cells | 74 | 2.04 | 3.39E-05 |
| Lipid and atherosclerosis | 190 | 1.78 | 4.13E-05 |
| Epstein-Barr virus infection | 185 | 1.79 | 4.59E-05 |
| Chemical carcinogenesis - reactive oxygen species | 208 | -1.77 | 5.14E-05 |
| Tight junction | 149 | 1.79 | 8.65E-05 |
| Thermogenesis | 219 | -1.71 | 1.43E-04 |
| IL-17 signaling pathway | 82 | 1.88 | 3.17E-04 |
| Estrogen signaling pathway | 106 | 1.82 | 3.17E-04 |
| Herpes simplex virus 1 infection | 159 | 1.71 | 3.30E-04 |
| Retrograde endocannabinoid signaling | 118 | -1.78 | 5.79E-04 |
| Shigellosis | 234 | 1.62 | 5.79E-04 |
| Metabolism of xenobiotics by cytochrome P450 | 56 | -2.00 | 5.84E-04 |
| Hepatitis B | 144 | 1.69 | 6.33E-04 |
| Apoptosis | 130 | 1.71 | 9.19E-04 |
| Proteasome | 44 | 1.93 | 1.02E-03 |
| Non-alcoholic fatty liver disease | 147 | -1.70 | 1.11E-03 |
| Drug metabolism - cytochrome P450 | 51 | -1.92 | 1.37E-03 |
| RIG-I-like receptor signaling pathway | 54 | 1.85 | 1.70E-03 |
| Chemical carcinogenesis - DNA adducts | 51 | -1.89 | 1.87E-03 |
| Neurotrophin signaling pathway | 113 | 1.70 | 2.09E-03 |
| Aminoacyl-tRNA biosynthesis | 44 | 1.87 | 2.22E-03 |

| | | | |
|---|---|---|---|
| Influenza A | 146 | 1.67 | 2.22E-03 |
| Endocytosis | 243 | 1.49 | 2.32E-03 |
| Chronic myeloid leukemia | 75 | 1.76 | 3.66E-03 |
| Fatty acid degradation | 39 | -1.86 | 5.32E-03 |
| Kaposi sarcoma-associated herpesvirus infection | 170 | 1.55 | 5.33E-03 |
| Spinocerebellar ataxia | 123 | 1.61 | 5.82E-03 |
| Acute myeloid leukemia | 63 | 1.73 | 6.36E-03 |
| Human papillomavirus infection | 289 | 1.46 | 6.51E-03 |
| IgSF CAM signaling | 262 | 1.45 | 6.73E-03 |
| Arginine biosynthesis | 19 | 1.89 | 6.86E-03 |
| Cytosolic DNA-sensing pathway | 68 | 1.70 | 6.88E-03 |
| Systemic lupus erythematosus | 119 | -1.59 | 8.02E-03 |
| Hippo signaling pathway | 139 | 1.59 | 8.02E-03 |
| Measles | 118 | 1.55 | 8.02E-03 |
| ATP-dependent chromatin remodeling | 111 | 1.58 | 9.97E-03 |
| Focal adhesion | 185 | 1.49 | 1.17E-02 |
| PD-L1 expression and PD-1 checkpoint pathway in cancer | 87 | 1.66 | 1.27E-02 |
| Adherens junction | 89 | 1.60 | 1.27E-02 |
| NF-kappa B signaling pathway | 103 | 1.56 | 1.27E-02 |
| Glycine, serine and threonine metabolism | 35 | -1.73 | 1.33E-02 |
| Gastric cancer | 124 | 1.53 | 1.36E-02 |
| Peroxisome | 78 | -1.55 | 1.39E-02 |
| Yersinia infection | 135 | 1.51 | 1.41E-02 |
| Mineral absorption | 48 | -1.66 | 1.62E-02 |
| Valine, leucine and isoleucine degradation | 47 | -1.69 | 1.70E-02 |
| Human cytomegalovirus infection | 201 | 1.44 | 1.70E-02 |
| Neuroactive ligand-receptor interaction | 204 | -1.43 | 1.91E-02 |
| Biosynthesis of nucleotide sugars | 36 | 1.71 | 2.02E-02 |
| B cell receptor signaling pathway | 81 | 1.56 | 2.03E-02 |
| Parkinson disease | 249 | -1.37 | 2.32E-02 |
| Leukocyte transendothelial migration | 99 | 1.52 | 2.41E-02 |
| Viral carcinogenesis | 191 | 1.42 | 2.41E-02 |
| T cell receptor signaling pathway | 114 | 1.51 | 2.44E-02 |
| Asthma | 22 | -1.77 | 2.56E-02 |
| Steroid biosynthesis | 19 | 1.75 | 2.56E-02 |
| Small cell lung cancer | 90 | 1.54 | 2.56E-02 |
| Toxoplasmosis | 105 | 1.52 | 2.56E-02 |
| Prion disease | 249 | -1.36 | 2.56E-02 |
| Amyotrophic lateral sclerosis | 328 | 1.33 | 2.56E-02 |
| Steroid hormone biosynthesis | 46 | -1.63 | 2.81E-02 |
| Endometrial cancer | 57 | 1.59 | 3.03E-02 |
| Arrhythmogenic right ventricular cardiomyopathy | 70 | 1.58 | 3.13E-02 |
| Renal cell carcinoma | 67 | 1.54 | 3.66E-02 |
| Thyroid cancer | 37 | 1.63 | 3.84E-02 |
| Cardiac muscle contraction | 68 | -1.50 | 3.92E-02 |
| Colorectal cancer | 85 | 1.48 | 4.02E-02 |
| Thyroid hormone synthesis | 63 | 1.54 | 4.43E-02 |
| Prostate cancer | 100 | 1.43 | 4.43E-02 |
| Necroptosis | 135 | 1.41 | 4.43E-02 |
| C-type lectin receptor signaling pathway | 95 | 1.40 | 4.43E-02 |
| Osteoclast differentiation | 129 | 1.39 | 4.51E-02 |
| Alanine, aspartate and glutamate metabolism | 32 | 1.61 | 4.61E-02 |
| JAK-STAT signaling pathway | 126 | 1.38 | 4.61E-02 |
| Huntington disease | 277 | -1.34 | 4.61E-02 |
| MAPK signaling pathway | 264 | 1.33 | 4.61E-02 |
| Integrated stress response (ISR) signaling pathway | 51 | 1.55 | 4.64E-02 |
| Chemical carcinogenesis - receptor activation | 165 | 1.38 | 4.78E-02 |
| MicroRNAs in cancer | 153 | 1.35 | 4.78E-02 |

[1]setSize: The number of genes in a given pathway that are represented in the analysis. [2]NES: Normalized enrichment score, reflecting the degree to which a gene set is overrepresented at the top (positive) or bottom (negative) of the ranked gene list, adjusted for gene set size.

**Table S5: Participant demographics of the single-cell RNAseq cohort**

| Characteristic | Overall<br>N = 13[1] | Long Covid<br>N = 9[1] | Recovered<br>N = 4[1] | p-value |
|---|---|---|---|---|
| **Age** (years) | 45 (34, 49) | 46 (43, 49) | 36 (32, 49) | 0.3 |
| **Sex** | | | | 0.9 |
| Female | 9 (69%) | 6 (67%) | 3 (75%) | |
| Male | 4 (31%) | 3 (33%) | 1 (25%) | |
| **Race/Ethnicity** | | | | 0.6 |
| American Indian or Alaska Native | 1 (7.7%) | 1 (11%) | 0 (0%) | |
| Asian | 2 (15%) | 2 (22%) | 0 (0%) | |
| Black/African American | 0 (0%) | 0 (0%) | 0 (0%) | |
| Hispanic/Latino | 1 (7.7%) | 0 (0%) | 1 (25%) | |
| White | 9 (69%) | 6 (67%) | 3 (75%) | |
| **Medical comorbidities** | | | | |
| Autoimmune disease | 3 (23%) | 2 (22%) | 1 (25%) | >0.9 |
| Cancer[2] | 0 (0%) | 0 (0%) | 0 (0%) | - |
| Diabetes | 0 (0%) | 0 (0%) | 0 (0%) | - |
| Hypertension | 1 (7.7%) | 1 (11%) | 0 (0%) | >0.9 |
| Lung disease[3] | 2 (15%) | 2 (22%) | 0 (0%) | >0.9 |
| **Body mass index** (kg/m2) | | | | 0.3 |
| <25 | 7 (54%) | 6 (67%) | 1 (25%) | |
| 25-29.9 | 5 (38%) | 2 (22%) | 3 (75%) | |
| >= 30 | 1 (7.7%) | 1 (11%) | 0 (0%) | |
| **Hospitalized for COVID-19** | 0 (0%) | 0 (0%) | 0 (0%) | |
| **First SVC2 infection date** | | | | >0.9 |
| March 2020 – April 2021 | 4 (31%) | 3 (33%) | 1 (25%) | |
| May 2021 – November 2021 | 0 (0%) | 0 (0%) | 0 (0%) | |
| >December 2021 | 9 (69%) | 6 (67%) | 3 (75%) | |
| **Time since first SCV2 infection** (days) | 507 (275, 674) | 507 (275, 674) | 499 (290, 1110) | 0.8 |
| **Time since most recent SCV2 infection** (days) | 439 (275, 636) | 439 (275, 515) | 499 (290, 1110) | 0.7 |
| **Vaccinated prior to colorectal biopsy** [4] | 13 (100%) | 9 (100%) | 4 (100%) | |
| **Quality of life by visual analog scale**[4] | 60 (50, 80) | 55 (50, 60) | 81 (80, 86) | 0.01 |
| **Long COVID symptom count**[4] | - | 8 (6, 8) | - | |
| **Symptoms reported**[5] | | | | |
| Fatigue | - | 7 (78%) | - | |
| Cardiopulmonary | - | 6 (67%) | - | |
| Gastrointestinal | - | 4 (44%) | - | |
| Smell/Taste disturbance | - | 1 (11%) | - | |
| Neurocognitive | - | 8 (89%) | - | |
| Post-Exertional Malaise | - | 3 (33%) | - | |
| Sleep disturbance | - | 5 (56%) | - | |

[1]n (%); Median (IQR). [2]Cancer requiring treatment within the 2 years before COVID-19. [3]Asthma, COPD, emphysema or bronchitis experienced in the 5 years prior to COVID-19. [4]At time of biopsy. [5]Cardiopulmonary: cough, shortness of breath, heart palpitations, chest pain. Gastrointestinal: diarrhea, nausea, loss of appetite, abdominal pain, vomiting, constipation. Neurocognitive: concentration problems, dizziness, balance problems, neuropathy, vision problems.

**Table S6: Participant demographics of the flow cytometry cohort**

| Characteristic | Overall N = 18[1] | Long Covid N = 10[1] | Recovered N = 8[1] | p-value |
|---|---|---|---|---|
| **Age** (years) | 46 (34, 56) | 47 (43, 54) | 37 (32, 58) | 0.5 |
| **Sex** | | | | 0.6 |
| Female | 12 (67%) | 6 (60%) | 6 (75%) | |
| Male | 6 (33%) | 4 (40%) | 2 (25%) | |
| **Race/Ethnicity** | | | | >0.9 |
| American Indian or Alaska Native | 1 (5.6%) | 1 (10%) | 0 (0%) | |
| Asian | 3 (17%) | 2 (20%) | 1 (13%) | |
| Black/African American | 0 (0%) | 0 (0%) | 0 (0%) | |
| Hispanic/Latino | 2 (11%) | 1 (10%) | 1 (13%) | |
| White | 12 (67%) | 6 (60%) | 6 (75%) | |
| **Medical comorbidities** | | | | |
| Autoimmune disease | 4 (22%) | 2 (20%) | 2 (25%) | >0.9 |
| Cancer[2] | 0 (0%) | 0 (0%) | 0 (0%) | - |
| Diabetes | 0 (0%) | 0 (0%) | 0 (0%) | - |
| Hypertension | 2 (11%) | 2 (20%) | 0 (0%) | 0.5 |
| Lung disease[3] | 2 (11%) | 2 (20%) | 0 (0%) | 0.5 |
| **Body mass index** (kg/m2) | | | | 0.8 |
| <25 | 10 (56%) | 6 (60%) | 4 (50%) | |
| 25-29.9 | 7 (39%) | 3 (30%) | 4 (50%) | |
| >= 30 | 1 (5.6%) | 1 (10%) | 0 (0%) | |
| **Hospitalized for COVID-19** | 1 (5.6%) | 1 (10%) | 0 (0%) | >0.9 |
| **First SVC2 infection era** | | | | 0.3 |
| March 2020 – April 2021 | 5 (28%) | 4 (40%) | 1 (13%) | |
| May 2021 – November 2021 | 0 (0%) | 0 (0%) | 0 (0%) | |
| >December 2021 | 13 (72%) | 6 (60%) | 7 (88%) | |
| **Time since first SCV2 infection** (days) | 536 (362, 729) | 511 (275, 729) | 597 (425, 764) | 0.5 |
| **Time since most recent SCV2 infection** (days) | 511 (361, 729) | 473 (275, 674) | 597 (424, 764) | 0.4 |
| **Vaccinated prior to colorectal biopsy**[4] | 18 (100%) | 10 (100%) | 8 (100%) | |
| **Quality of life by visual analog scale**[4] | 73 (55, 80) | 58 (50, 65) | 81 (80, 88) | <0.001 |
| **Long COVID symptom count**[4] | - | 8 (4, 8) | - | |
| **Symptoms reported**[5] | | | | |
| Fatigue | - | 7 (70%) | - | |
| Cardiopulmonary | - | 6 (60%) | - | |
| Gastrointestinal | - | 4 (40%) | - | |
| Smell/Taste disturbance | - | 1 (10%) | - | |
| Neurocognitive | - | 9 (90%) | - | |
| Post-Exertional Malaise | - | 3 (30%) | - | |
| Sleep disturbance | - | 6 (60%) | - | |

[1]n (%); Median (IQR). [2]Cancer requiring treatment within the 2 years before COVID-19. [3]Asthma, COPD, emphysema or bronchitis experienced in the 5 years prior to COVID-19. [4]At time of biopsy. [5]Cardiopulmonary: cough, shortness of breath, heart palpitations, chest pain. Gastrointestinal: diarrhea, nausea, loss of appetite, abdominal pain, vomiting, constipation. Neurocognitive: concentration problems, dizziness, balance problems, neuropathy, vision problems. Symptom data were unavailable for one participant

**Table S7: Participant demographics of the Xenium cohort**

| Characteristic | Overall N = 6[1] | Long Covid N = 4[1] | Recovered N = 2[1] | p-value |
|---|---|---|---|---|
| **Age** (years) | 48 (33, 61) | 58 (48, 64) | 32 (30, 33) | 0.13 |
| **Sex** | | | | >0.9 |
| Female | 3 (50%) | 2 (50%) | 1 (50%) | |
| Male | 3 (50%) | 2 (50%) | 1 (50%) | |
| **Race/Ethnicity** | | | | 0.9 |
| American Indian or Alaska Native | 1 (17%) | 1 (25%) | 0 (0%) | |
| Asian | 0 (0%) | 0 (0%) | 0 (0%) | |
| Black/African American | 0 (0%) | 0 (0%) | 0 (0%) | |
| Hispanic/Latino | 2 (33%) | 1 (25%) | 1 (50%) | |
| White | 3 (50%) | 2 (50%) | 1 (50%) | |
| **Medical comorbidities** | | | | |
| Autoimmune disease | 0 (0%) | 0 (%) | 0 (0%) | - |
| Cancer[2] | 0 (0%) | 0 (0%) | 0 (0%) | - |
| Diabetes | 1 (17%) | 1 (25%) | 0 (0%) | >0.9 |
| Hypertension | 0 (0%) | 0 (0%) | 0 (0%) | - |
| Lung disease[3] | 1 (17%) | 1 (25%) | 0 (0%) | >0.9 |
| **Body mass index** (kg/m2) | | | | 0.6 |
| <25 | 1 (17%) | 0 (0%) | 1 (50%) | |
| 25-29.9 | 4 (67%) | 3 (75%) | 1 (50%) | |
| >= 30 | 1 (17%) | 1 (25%) | 0 (0%) | |
| **Hospitalized for COVID-19** | 1 (17%) | 1 (25%) | 0 (0%) | >0.9 |
| **First SVC2 infection era** | | | | 0.5 |
| March 2020 – April 2021 | 2 (33%) | 2 (50%) | 0 (0%) | |
| May 2021 – November 2021 | 0 (0%) | 0 (0%) | 0 (0%) | |
| >December 2021 | 4 (67%) | 2 (50%) | 2 (100%) | |
| **Time since first SCV2 infection** (days) | 356 (211, 636) | 353 (200, 584) | 427 (218, 636) | 0.8 |
| **Time since most recent SCV2 infection** (days) | 215 (188, 636) | 200 (157, 443) | 427 (218, 636) | 0.5 |
| **Vaccinated prior to colorectal biopsy**[4] | 5 (83%) | 3 (75%) | 2 (100%) | |
| **Quality of life by visual analog scale**[4] | 5 (83%) | 3 (75%) | 2 (100%) | |
| **Long COVID symptom count**[4] | | | | |
| **Symptoms reported**[5] | - | 4 (2, 8) | - | |
| Fatigue | 80 (70, 80) | 70 (60, 80) | 85 (80, 90) | 0.2 |
| Cardiopulmonary | - | 4 (100%) | - | |
| Gastrointestinal | - | 2 (50%) | - | |
| Smell/Taste disturbance | - | 2 (50%) | - | |
| Neurocognitive | - | 1 (25%) | - | |
| Post-Exertional Malaise | - | 2 (50%) | - | |
| Sleep disturbance | - | 1 (25%) | - | |

[1]n (%); Median (IQR). [2]Cancer requiring treatment within the 2 years before COVID-19. [3]Asthma, COPD, emphysema or bronchitis experienced in the 5 years prior to COVID-19. [4]At time of biopsy. [5]Data missing for one participant.[6]Cardiopulmonary: cough, shortness of breath, heart palpitations, chest pain. Gastrointestinal: diarrhea, nausea, loss of appetite, abdominal pain, vomiting, constipation. Neurocognitive: concentration problems, dizziness, balance problems, neuropathy, vision problems.

**Table S8: List of all 477 genes included in the Xenium panel**

| | | | | | | | | |
|---|---|---|---|---|---|---|---|---|
| ABCC11 | C7 | CDK1 | DPP6 | GPRC5A | KLRC3 | MYLK | S100A12 | CMV_UL111A_vIL10 |
| ACE2 | CA4 | CDK15 | DPT | GPX2 | KLRD1 | MZB1 | S100B | CMV_UL83_pp65 |
| ACKR1 | CAPN8 | CENPF | DST | GYPA | KLRK1 | NAT8 | SCGB2A1 | EBV_BCRF1_vIL10 |
| ACTA2 | CARD8 | CFAP53 | DUSP2 | GYPB | KNG1 | NCAM1 | SCGN | HIV_Gag_unspliced |
| ACTG2 | CASP1 | CFB | EBV_BILF1 | GZMA | KRT20 | NFKB1 | SELE | HIV_Pol_unspliced |
| ADAM17 | CAV1 | CFHR1 | EBV_EBNA1 | GZMB | KRT7 | NKG7 | SELL | HIV_Tat-RevIII |
| ADAM28 | CAVIN1 | CFHR3 | ECSCR | GZMK | LAG3 | NLRP3 | SEMA3C | HIV_Tat-RevII_spliced |
| ADAMTS1 | CAVIN2 | CFTR | EDN1 | HAMP | LAMP3 | NOS2 | SERPINB2 | HIV_Tat_RevI |
| ADGRE1 | CCDC39 | CHGA | EDNRB | HAVCR2 | LCK | NPDC1 | SERPINB3 | SARS-CoV-2_E |
| ADGRL4 | CCDC78 | CLCA1 | EGFL7 | HEMGN | LGI4 | NTN4 | SERPINB9 | SARS-CoV-2_E_subgenomic |
| ADH1C | CCL19 | CLCA2 | EGFR | HEPACAM2 | LGR5 | OGN | SFRP2 | SARS-CoV-2_N |
| ADH4 | CCL20 | CLEC10A | EHF | HES4 | LIF | OPRPN | SFRP4 | SARS-CoV-2_ORF10 |
| ADIPOQ | CCL27 | CLEC14A | ELF5 | HIGD1B | LILRA4 | PCNA | SFTA2 | SARS-CoV-2_ORF3A |
| AGER | CCL5 | CLEC4E | EPCAM | HIV_LTR | LILRA5 | PCOLCE | SH2D3C | SARS-CoV-2_ORF8 |
| AGR3 | CCNB2 | CLEC9A | ERBB2 | HIV_Nef | LILRB2 | PCP4 | SLAMF1 | SARS-CoV-2_RdRp |
| AHSP | CCR2 | CLECL1 | ERG | HIV_PolyA | LILRB4 | PCSK2 | SLAMF7 | SARS-CoV-2_S |
| AIF1 | CCR7 | CLIC6 | ESR1 | HIV_TAR | LPL | PDCD1 | SLC18A2 | SARS-CoV-2_S_antisense |
| AKT1 | CD14 | CNN1 | FABP3 | HLA-DQB2 | LTBP2 | PDGFRA | SLC22A8 | THBS2 |
| ALAS2 | CD163 | COCH | FAS | HMGCS2 | LY6D | PDGFRB | SLC26A2 | THY1 |
| ALDH1A3 | CD177 | COL17A1 | FBLN1 | HPGDS | LY86 | PDPN | SLC26A3 | TIGIT |
| AMY2A | CD19 | COL5A2 | FBN1 | HPX | LYVE1 | PEBP4 | SLC4A1 | TIMP3 |
| ANGPT2 | CD1A | COL8A1 | FCER1A | ICOS | MAF | PECAM1 | SMIM24 | TIMP4 |
| ANPEP | CD1C | CPA3 | FCGR1A | IFNA1 | MALL | PGR | SMYD2 | TM4SF18 |
| APCDD1 | CD1E | CRHBP | FCGR3A | IFNAR1 | MAMDC2 | PIK3CD | SNAI1 | TM4SF4 |
| APOA5 | CD2 | CRISPLD2 | FCGR3B | IFNB1 | MAP1LC3B | PLA2G7 | SNCA | TMC5 |
| APOBEC3A | CD24 | CSF2RA | FCN1 | IFNG | MARCO | PLAC9 | SNCG | TMEM100 |
| APOLD1 | CD247 | CSF3 | FCN2 | IFNGR1 | MCEMP1 | PLCG2 | SNTN | TMEM174 |
| AQP2 | CD27 | CTLA4 | FGFBP1 | IGF1 | MCF2L | PLD4 | SOX17 | TMEM52B |
| AQP3 | CD274 | CTSG | FGFBP2 | IGSF6 | MDM2 | PLIN4 | SOX18 | TMPRSS2 |
| AQP8 | CD28 | CTSK | FGFR4 | IL10 | MEDAG | PMP22 | SOX2 | TNC |
| AQP9 | CD300E | CTSL | FGL2 | IL18 | MEF2C | PPARG | SPDEF | TNFAIP3 |
| AR | CD34 | CXCL10 | FHL2 | IL1B | MEST | PPP1R1A | SPI1 | TNFRSF13B |
| ARFGEF3 | CD38 | CXCL2 | FKBP11 | IL1R2 | MET | PPP1R1B | SPIB | TNFRSF17 |
| ASCL1 | CD3D | CXCL5 | FOXA1 | IL1RAPL1 | MFAP5 | PPY | SRPX | TNFRSF18 |
| ASCL3 | CD3E | CXCL6 | FOXI1 | IL1RL1 | MKI67 | PRDM1 | SST | TNFRSF25 |
| ASPN | CD4 | CXCL8 | FOXJ1 | IL22 | MLANA | PRF1 | STAT4 | TNFRSF8 |
| ATG5 | CD40 | CXCL9 | FOXP3 | IL2RA | MLPH | PRG4 | STAT5A | TNFRSF9 |
| ATG7 | CD40LG | CXCR4 | FSTL3 | IL3RA | MMP12 | PROX1 | STAT5B | TNFSF13B |
| BAMBI | CD5 | CXCR5 | FXYD2 | IL4I1 | MMP9 | PTGDS | STC1 | TNFSF8 |
| BANK1 | CD5L | CYP1A1 | GATA2 | IL6 | MMRN1 | PTN | STC2 | TOP2A |
| BASP1 | CD6 | CYP2A7 | GATM | IL7R | MMRN2 | PTPRC | STEAP4 | TRAC |
| BBOX1 | CD68 | CYP2B6 | GCG | INMT | MNDA | PVALB | TAC1 | TREM2 |
| BCL2L11 | CD69 | CYP2F1 | GDF15 | INS | MPEG1 | RAMP2 | TAT | TRPM2 |
| BECN1 | CD7 | CYP3A4 | GEM | IQGAP2 | MRC1 | RAPGEF3 | TBX3 | TSPAN19 |
| BMX | CD70 | CYP4B1 | GHRL | IRF8 | MS4A1 | RBP5 | TCF15 | UBE2C |
| BTNL9 | CD79A | CYTIP | GKN2 | ITGAM | MS4A2 | RERGL | TCF4 | UMOD |
| C15orf48 | CD80 | DERL3 | GLIPR1 | ITK | MS4A4A | RETN | TCF7 | UPK3B |
| C1QB | CD83 | DES | GLYATL1 | KCNK3 | MS4A6A | RGS16 | TCIM | VCAN |
| C1orf162 | CD86 | DIRAS3 | GNG11 | KCNMA1 | MTOR | RIDA | TCL1A | VSIG4 |
| C1orf194 | CD8A | DMBT1 | GNLY | KIT | MTRNR2L11 | RND1 | TENT5C | VWA5A |
| C20orf85 | CD8B | DNAAF1 | GPC1 | KLK11 | MYBPC1 | RPS6 | TFF2 | VWF |
| C5orf46 | CD93 | DNASE1L3 | GPC3 | KLRB1 | MYC | RTKN2 | TFPI | WNT2 |
| C6orf118 | CDH16 | DPEP1 | GPR183 | KLRC1 | MYH11 | S100A1 | THAP2 | XCL2 |

**Table S9: Flow cytometry reagents**

| | Color | Target | Company | Catalog ID | Clone | Raised in | Lot ID | Dilution |
|---|---|---|---|---|---|---|---|---|
| **UV Laser** | BUV395 | streptavidin | BD | 564176 | - | - | 5076912 | 50 |
| | | IgA-biotin | Southernbio | 2052-08 | - | Goat F(ab')2 | 0521-Z244Z | 500 |
| | Zombie UV | LD | Biolegend | 423107 | - | - | B334687 | 1000 |
| **Violet Laser** | BV421 | CCR7 | Biolegend | B360659 | G043H7 | Mu | B360659 | 30 |
| | cFV450 | CD45RA | Cytek | R7-20122 | HI100 | Mu | F-022723-02 | 30 |
| | BV510 | IgM | Biolegend | B365834 | MHM-88 | Mu | B365834 | 150 |
| | cFV547 | CD20 | Cytek | R7-20112 | 2H7 | Mu | F-032823-04 | 30 |
| | BV570 | CD3 | Biolegend | B360619 | UCHT1 | Mu | B360619 | 30 |
| | BV650 | CD28 | Biolegend | B362788 | CD28.2 | Mu | B362788 | 30 |
| | BV711 | CD38 | Biolegend | B360789 | HIT2 | Mu | B360789 | 30 |
| | BV750 | CD56 | Biolegend | B362796 | 5.1H11 | Mu | B362796 | 30 |
| | BV785 | PD-1 | Biolegend | B360654 | EH12.2H7 | Mu | B360654 | 30 |
| **Blue Laser** | cFB515 | CD141 | Cytek | R7-20114 | M80 | Mu | F-080822-04 | 200 |
| | cFB532 | CD8 | Cytek | R7-20124 | SK1 | Mu | F-121922-01 | 150 |
| | cFB548 | CD14 | Cytek | R7-20116 | 63D3 | Mu | F-032123-01 | 300 |
| | cFB690 | HLA-DR | Cytek | R7-20126 | L243 | Mu | F-081122-05 | 150 |
| **Yellow Laser** | cFBYG575 | CD25 | Cytek | R7-20128 | BC96 | Mu | F-122222-01 | 30 |
| | cFYG584 | CD4 | Cytek | R7-20042 | SK3 | Mu | F-012022-01 | 30 |
| | cFBYG610 | CD16 | Cytek | R7-20130 | 3G8 | Mu | F-021723-02 | 150 |
| | cFBYG667 | IgD | Cytek | R7-20138 | IA6-2 | Mu | F-121721-02 | 200 |
| | cFBYG710 | TCRgd | Cytek | R7-20136 | B1 | Mu | F-021422-01 | 30 |
| | cFBYG781 | CD11c | Cytek | B457720 | 3.9 | Mu | B457720 | 30 |
| **Red Laser** | cFR659 | CD127 | Cytek | R7-20078 | A019D5 | Mu | F-013023-01 | 75 |
| | cFR668 | CD1c | Cytek | R7-20120 | L161 | Mu | F-010923-03 | 300 |
| | cFR685 | CD19 | Cytek | R7-20118 | HIB19 | Mu | F-081222-03 | 30 |
| | cFR720 | CD123 | Cytek | R7-20014 | 6H6 | Mu | F-102521-03 | 30 |
| | cFR780 | CD45 | Cytek | R7-20134 | 2D1 | Mu | F-020322-01 | 30 |
| | cFR840 | CD27 | Cytek | R7-20082 | QA17A18 | Mu | F-120922-01 | 30 |

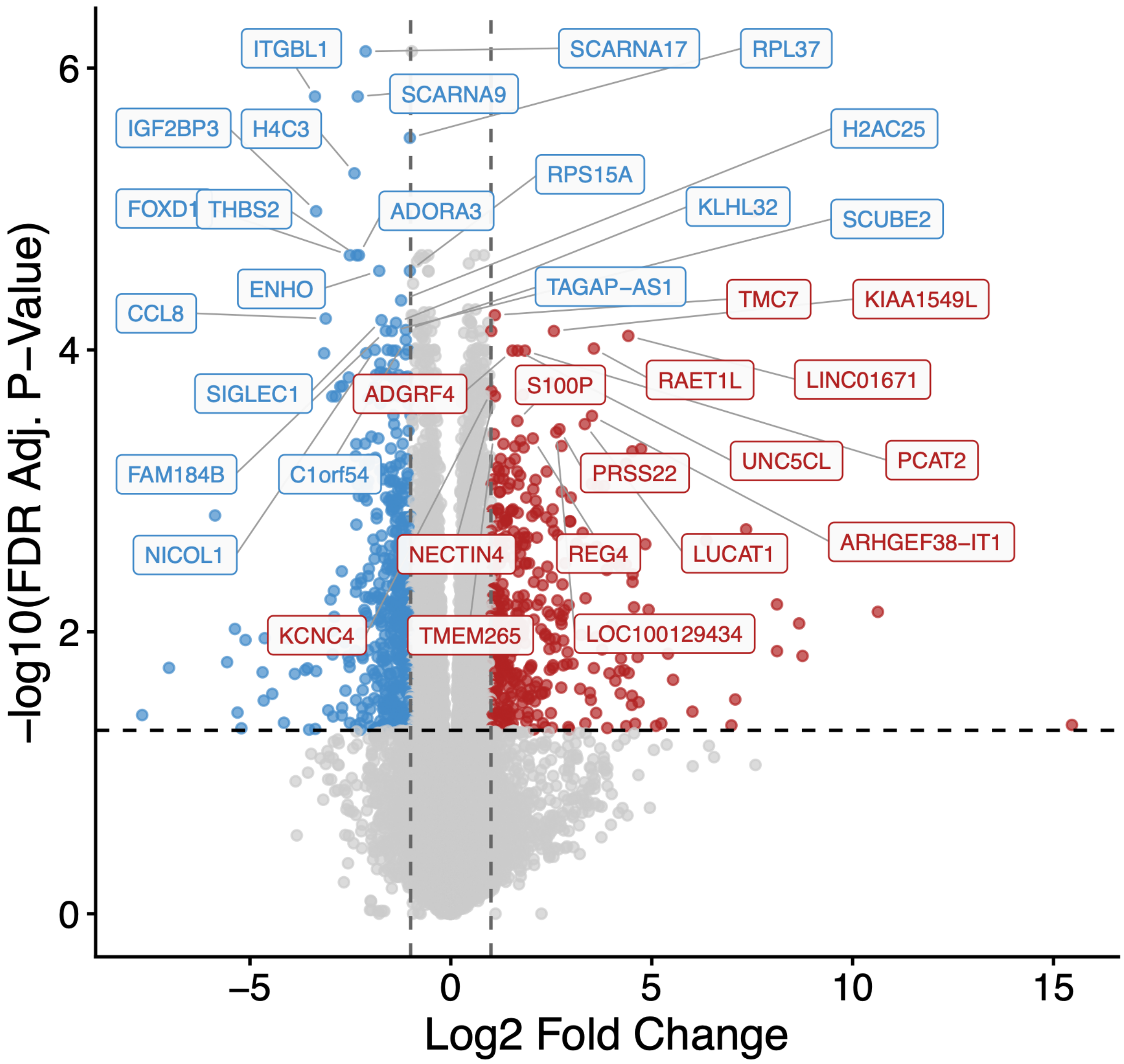

**Figure S1**: Volcano plots displaying differential expression analysis from colorectal bulk tissue whole-transcriptome polyA-RNAseq in people with LC versus recovered controls is shown. Significant DEGs (after FDR-adjustment) represented by red (upregulated) and blue (downregulated) dots. Top DEGs are labeled.

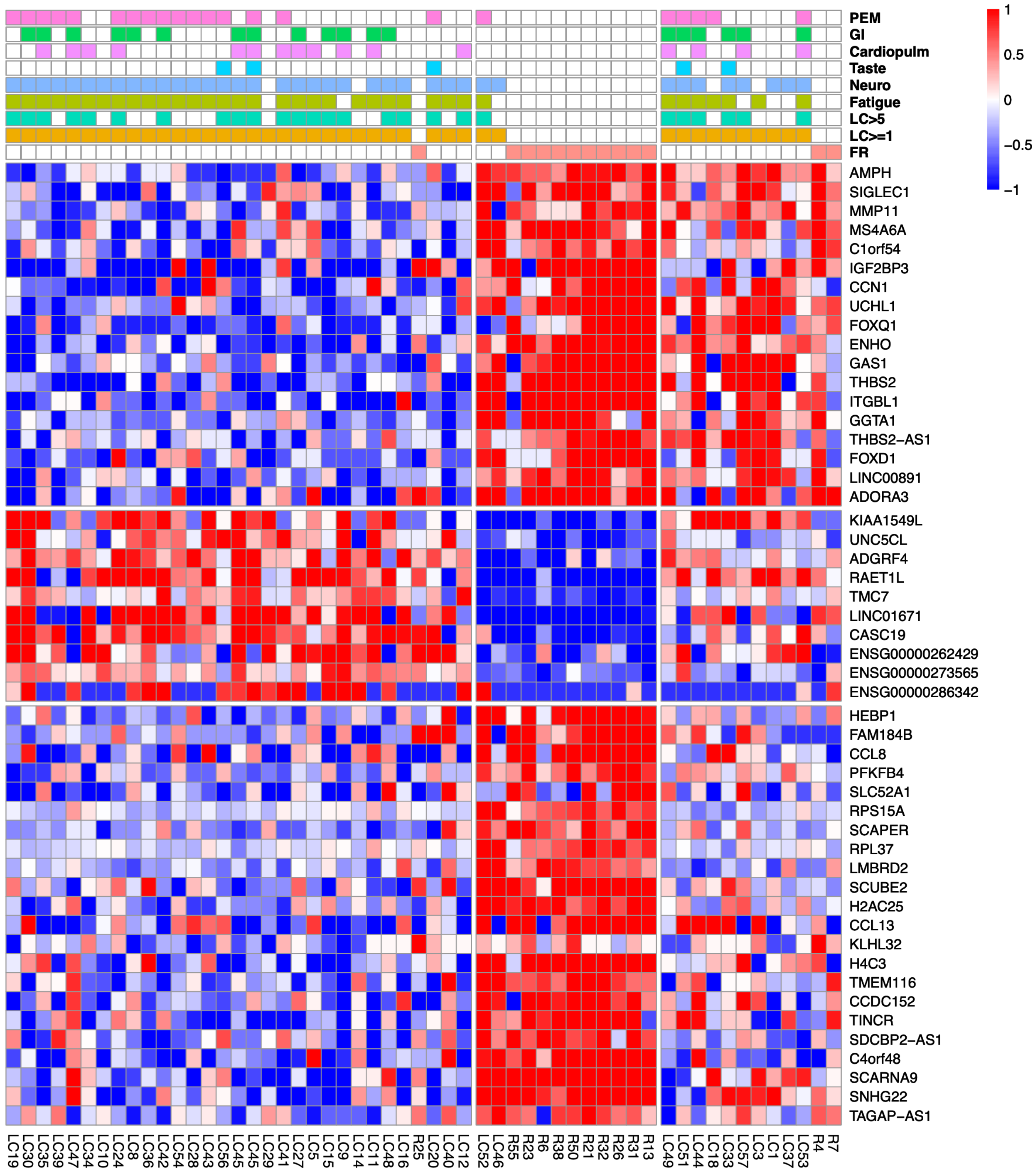

**Figure S2**: A heat map of the top 50 FDR-adjusted DEGs from colorectal bulk tissue whole-transcriptome polyA-RNAseq in people with LC versus recovered controls. The top horizontal bars represent which participants experienced each LC symptom phenotype at the time of tissue sampling (PEM = post exertional malaise; GI = gastrointestinal, taste = smell and taste disturbances; fatigue; neuro = neurocognitive symptoms; LC>5 = participants experiencing greater than 5 LC symptoms, and LC>1 participants experiencing one or more LC symptom). A complete list of all 4757 significant DEGs shown in Supplemental Data File S1.

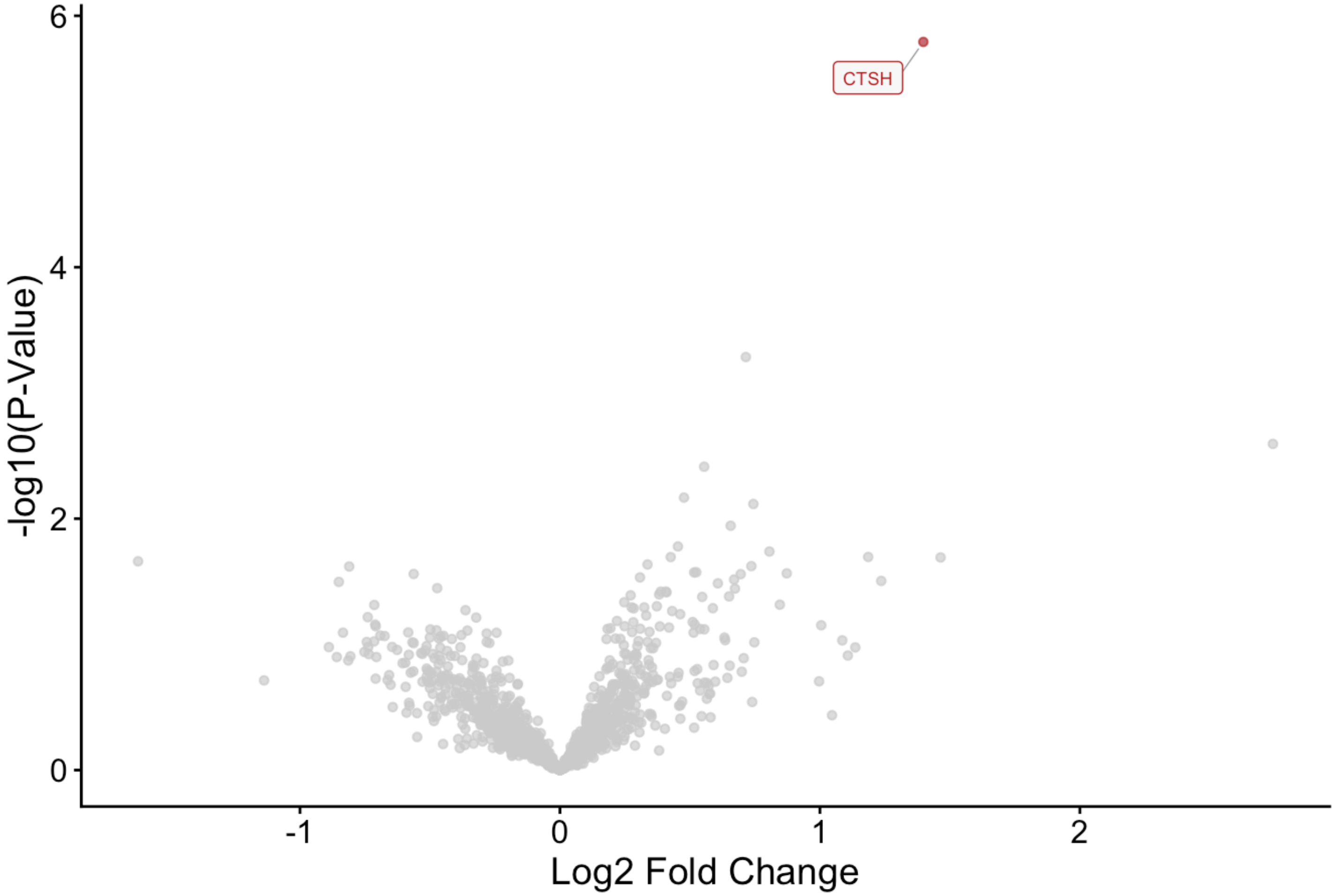

**Figure S3**: Volcano plot displaying differential expression analysis of deep plasma proteomic (Olink Reveal) data in people with LC experiencing taste and smell disturbances vs recovered participants.

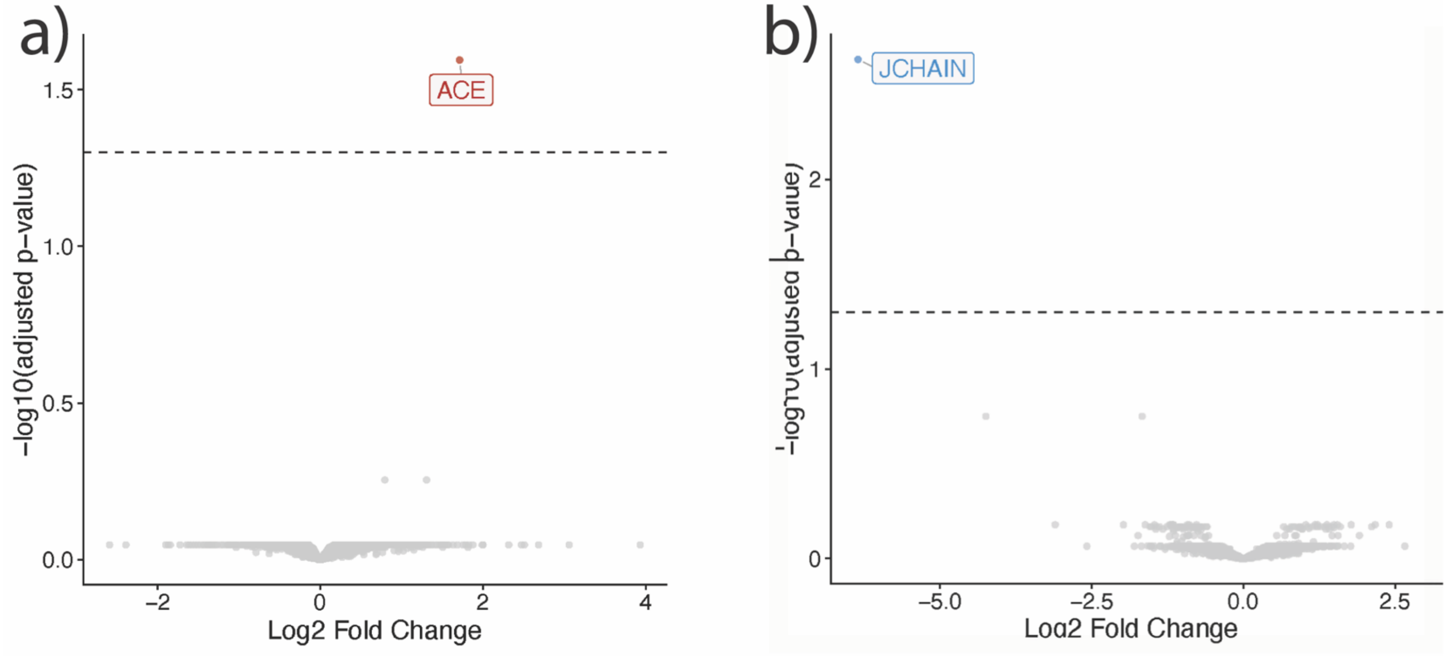

**Figure S4**: Volcano plots displaying differential expression analysis from unbiased single-cell transcriptomics comparing participants with LC to recovered controls. Significant Differentially Expressed Genes (DEGs) (after FDR-adjustment) represented by red (upregulated) and blue (downregulated) dots.

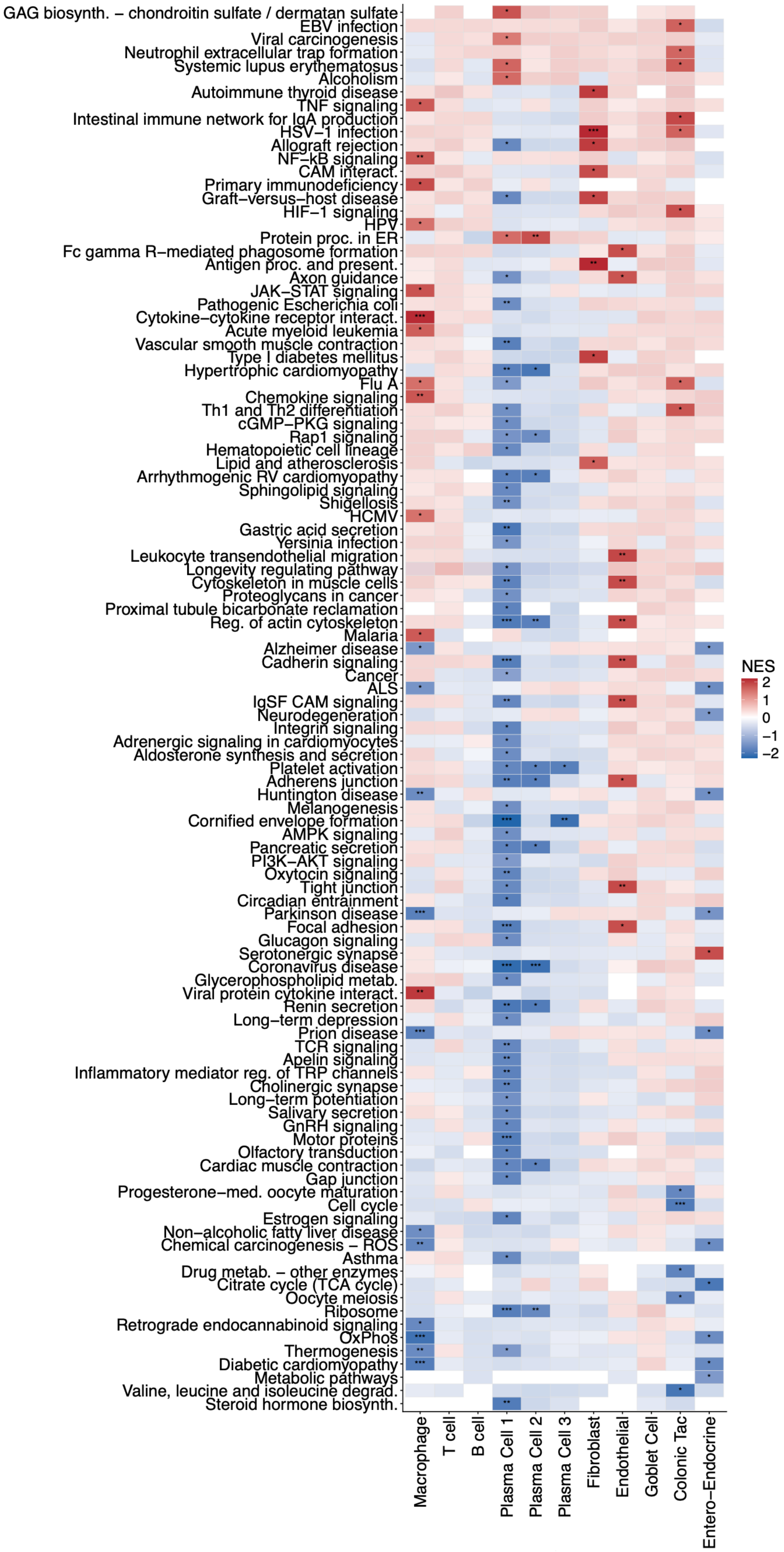

**Fig S5**: KEGG Gene set enrichment analyses from unbiased single-cell transcriptomics comparing participants with LC to recovered controls. Significant gene expression pathways for each unique cell cluster are summarized, shaded by Normalized Enrichment Scores (NES; upregulated in LC = red, downregulate in LC = blue). Significant different NES from FDR-adjusted analyses are labeled with asterisks (*= 0.05, ** = 0.01. *** = 0.001, etc.). Significant pathways are opaque while nonsignificant pathways are transparent.

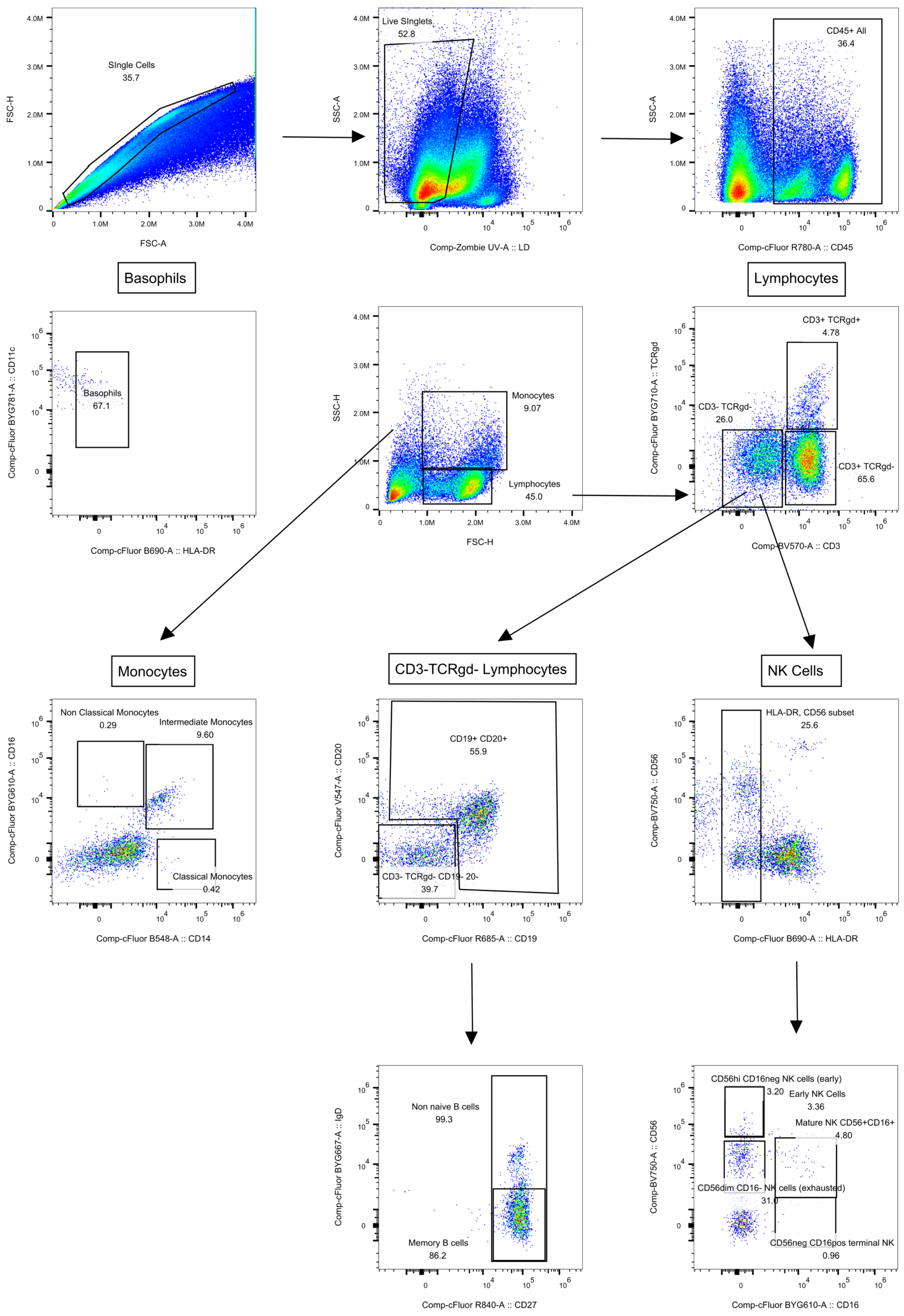

**Fig S6**: Flow cytometry anchor cell gating strategy

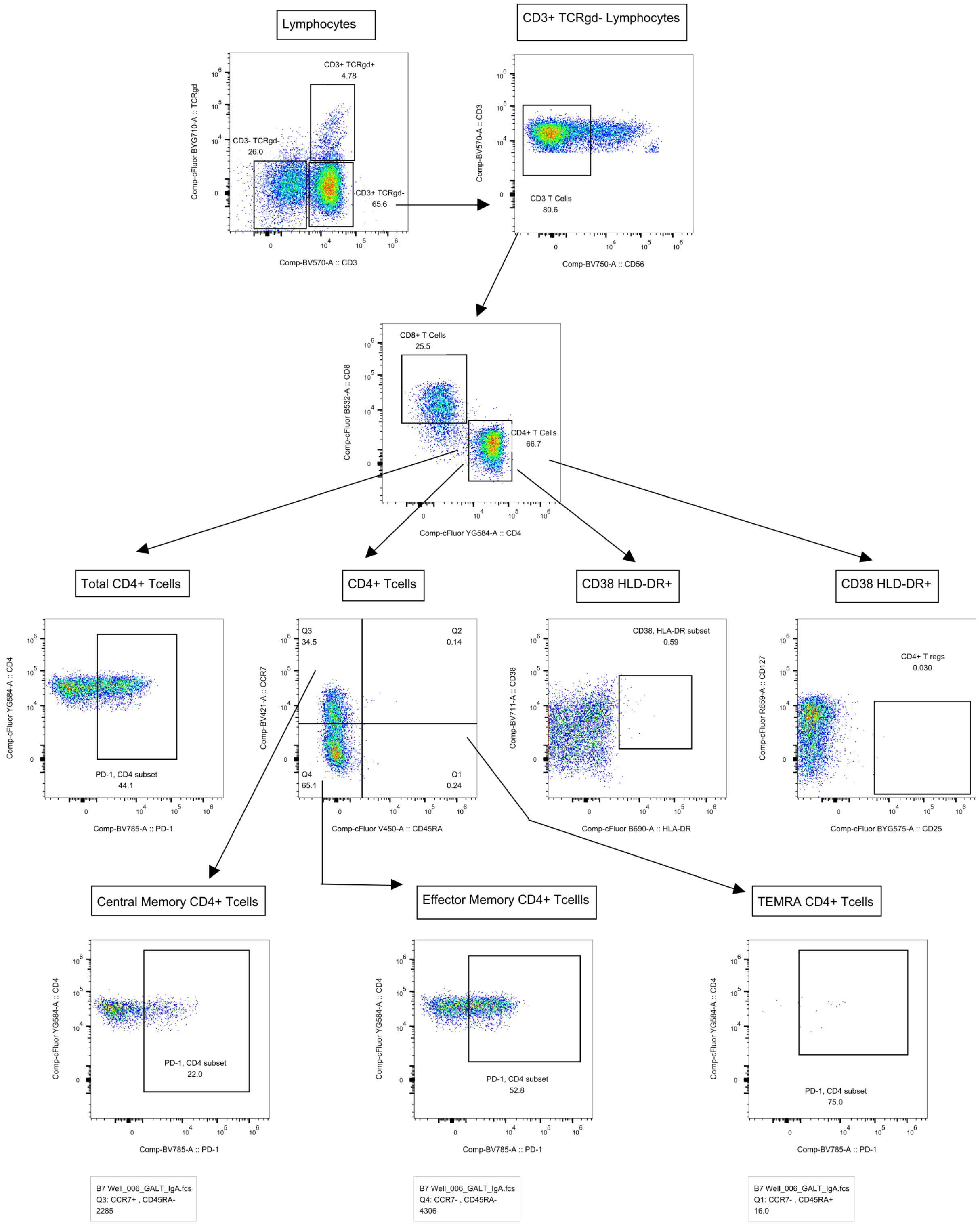

**Fig S7**: Flow cytometry CD4+ cell gating strategy

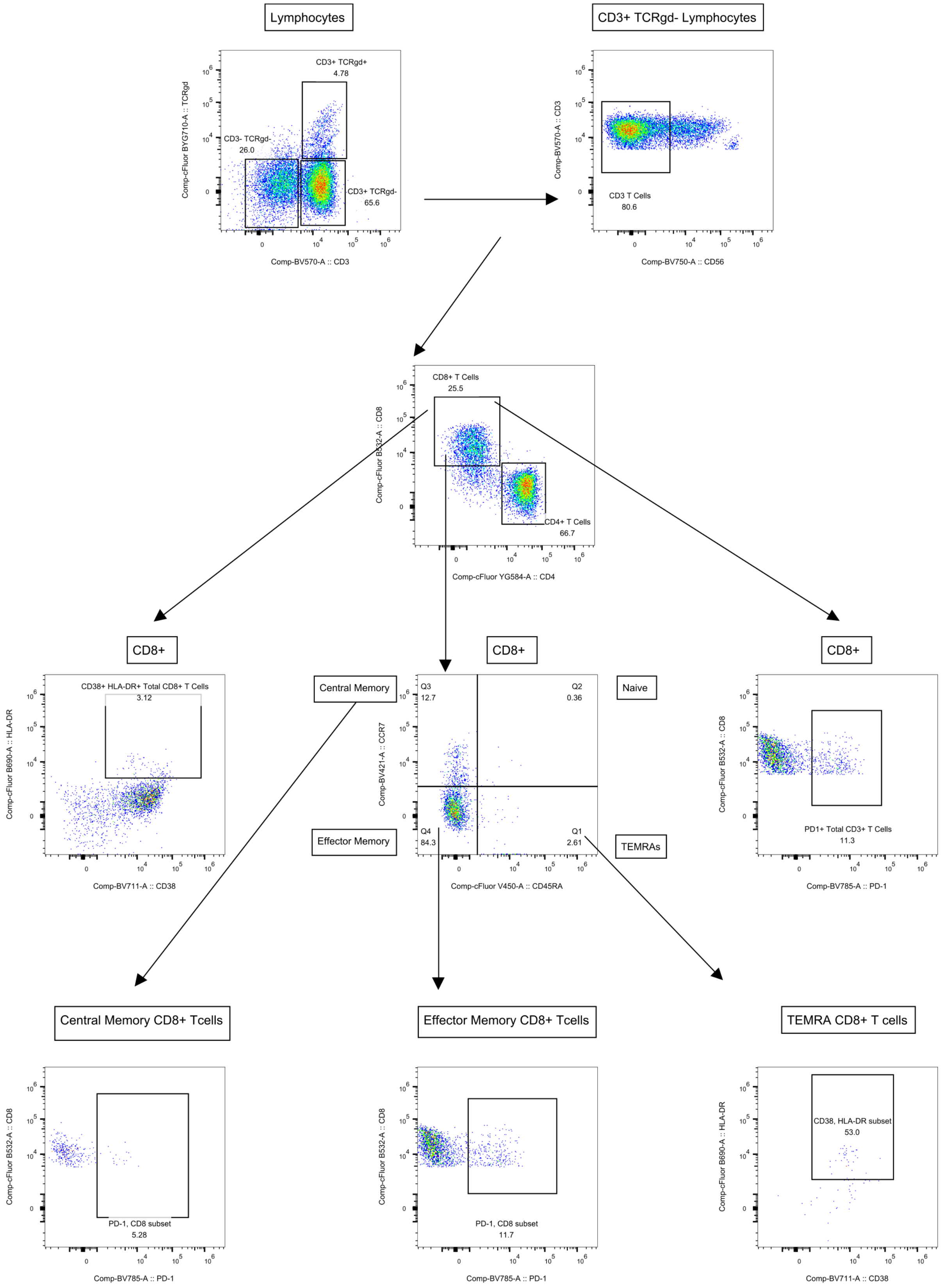

**Fig S8**: Flow cytometry CD8+ cell gating strategy

**Data file S1:** Significant DEGs from bulk RNAseq analysis.
